# Trophic rewilding pumas and guanacos: estimating the potential to enhance carbon sequestration in a Patagonian grassland ecosystem

**DOI:** 10.64898/2026.07.28.740956

**Authors:** Matteo Rizzuto, Ingrid Espinoza, Cristián Saucedo, Oswald J. Schmitz

**Author notes:** Department of Biology, Concordia University, Montreal, Quebec, Canada; Department of Biology, Memorial University of Newfoundland, St. John’s, NL, Canada.

## Abstract

1. Trophic rewilding, the practice of restoring animal species to recover lost ecosystem functions, has been proposed as a nature-based climate change solution (NbCS) due to animal biogeochemical influences potentially extending to ecosystem carbon capture.
2. We examined this potential using a case study of puma (*Puma concolor*) and guanaco (*Lama guanicoe*) restoration in a grassland ecosystem in Patagonia National Park, Chile. We parameterized a model of animal-driven ecosystem carbon dynamics with published measurements from Patagonian grasslands and compared three scenarios: (1) a no-rewilding baseline; (2) rewilding only guanaco; and (3) rewilding guanaco and puma. We estimated net primary productivity (NPP), net ecosystem carbon balance (NECB), and plant and soil carbon stocks. Using differences among scenarios, we estimated ecosystem carbon gains attributable to rewilding guanacos and pumas, and validated baseline estimates using published carbon capture data for Patagonian grasslands.
3. Patagonian grasslands with pumas and guanacos could capture (NPP and NECB) 1.27–2.5 times more carbon, and increase plant carbon by 1.76–3.25 times, above the no-rewilding baseline. Large uncertainties in parameter values make estimating soil carbon challenging, but the rewilded ecosystem could store up to 1.16 times more soil carbon, or up to 0.57 times less.
4. The model estimates that NECB in the rewilded ecosystem could amount to 94.41 t C km^-2^ y^-1^ (94.38 t C km^-2^ y^-1^–94.44 t C km^-2^ y^-1^) of which 23%–43% attributable to animal effects. Plant carbon storage estimates were ∼290 t C km^-2^ (280–300 t C km^-2^), of which 43%–67% attributable to animal effects. Finally, the model estimated soil C stock gains up to 178 t C km^-2^, or losses up to 4300 t C km^-2^.
5. *Practical implications.* We illustrate how to develop first approximation estimates of carbon capture and storage to help assess the feasibility of trophic rewilding as a NbCS. Our modelling revealed that restoring a puma-guanaco trophic cascade could be a feasible NbCS, and identified looming uncertainties about the fate of carbon that need further empirical exploration. More generally, the modelling identifies key measurements that can inform whether restoring trophic cascades can contribute to NbCS.

## 1 INTRODUCTION

Trophic rewilding is a restoration strategy whereby animal species are reintroduced to landscapes they once occupied to restore their functional roles within ecosystems (Svenning et al., 2019; 2024). One key functional role of animals is the top-down control they exert via trophic cascades (Svenning et al., 2019; 2024): top predators change their herbivore prey abundance and foraging, altering herbivore impacts on plants’ abundance and diversity (Terborgh et al., 2010) and potentially instigating further changes to photosynthesis and thus ecosystem carbon cycling (Schmitz et al., 2018). Evidence from experimental predator manipulations at local spatial scales (Strickland et al., 2012; Wyatt et al., 2021) and empirical analyses of relationships between top predator occurrence and carbon capture and storage across broad spatial extents (Schmitz et al., 2017; Roberts et al. 2025) provide proof-of-concept for this principle. These findings suggest that trophic rewilding could be a promising nature-based solution (Schmitz et al., 2023).

However, translating this promise into actionable nature-based climate solutions (NbCS) for particular species and ecosystems requires addressing important unknowns. Foremost, it remains generally unknown whether, where, and by how much trophic cascades will enhance carbon capture and storage in an ecosystem, because few studies have quantified the net effects of trophic cascades on whole ecosystem carbon cycling at landscape scales. Existing evidence indicates that trophic interactions between a given predator species and prey vary with ecosystem type, leading to either positive or negative effects on carbon capture and storage (Wilmers and Schmitz, 2016; Roberts et al., 2025).

These unknowns loom large at a time when there is a recognized, growing need to rewild ecosystems with large predators and their prey to restore ecosystem intactness and functionality, including functions that could enhance resilience to climate change (Svenning et al., 2019; 2024; Roberts et al., 2025; Spracklen et al. 2025). Yet, conducting preliminary field studies that address these unknowns to justify implementing trophic rewilding as a NbCS is challenging. Obtaining robust measures of carbon capture and storage requires spatially extensive, lengthy, and costly research that assesses carbon budgets between locations with and without focal animals (e.g., Naidu et al., 2022; Roberts et al., 2025). Moreover, the absence of focal animal species within candidate ecosystems for rewilding often precludes undertaking such preliminary assessments.

Ecosystem modelling offers an alternative, preliminary approach to address these unknowns and account for animal effects on ecosystem carbon capture and storage (Rizzuto et al. 2024; Spracklen et al. 2025). When parametrized using data from published studies, such models can be used to systematically explore scenarios that estimate and compare ecosystem carbon budgets with different combinations of absent vs present animals. Such analyses can provide insight into how much carbon capture and storage could be attributable to the functional role of each target predator and prey species (Spracklen et al. 2025). If modelling results demonstrate feasibility, they further provide first-approximation quantitative estimates of animal impacts that can be verified using trophic rewilding as an experiment. This would involve monitoring changes in carbon budgets of focal systems before, during, and after animal populations are rewilded, including using animal exclusions as controls (e.g., Kaštovská et al., 2024).

Here we report on such modelling to assess whether restoring a trophic cascade could become a NbCS. We examine a case study where the once-present apex predator puma (*Puma concolor*) and the once-dominant herbivore guanaco (*Lama guanicoe*) are rewilded into a grassland ecosystem in Patagonia National Park (PNP), Chile. The modeling produces estimates that fall within the range of literature-based empirical measurements of carbon capture and storage, suggesting that trophic rewilding pumas and guanacos could be a feasible NbCS in this grassland system. However, the model estimates are sensitive to changes in the magnitude of certain parameters related to puma and guanaco impacts on biogeochemical processes, for which accurate measures are lacking. Thus, the modeling highlights crucial new empirical research needed in this system, and in wildlife ecology more generally, to better assess whether the functional impact of trophic cascades on ecosystem carbon budgets will be sufficient to promote trophic rewilding as an NbCS.

## 2 METHODS

### 2.1 The focal ecosystem

The focal landscape is the Valle Chacabuco, Aysén district, Chile (Figure 1). Guanaco was the dominant grazer, and puma the once-dominant predator. However, both species’ abundances declined after the introduction of livestock to the landscape—either to control grazing competition (Hernandez et al., 2019) or control predation on livestock (Donadio et al. 2022). Recently, parts of the region were returned to natural conditions after century-long livestock ranching through the creation of PNP in 2018. PNP was established by Tompkins Conservation, after acquiring 83,000 ha of land in Estancia Valle Chacabuco. After these lands were donated to the Chilean State, two adjacent National Reserves were incorporated into PNP, expanding it to > 300,000 ha (3,000 km²) and subsequently unified into a single national park (Tompkins, 2024). We focus on the 485.91 km^2^ subregion of PNP within the Valle Chacabuco comprising habitats that support grazing, including coironales (Patagonian grassland steppe), perennial grassland, vegas (wet meadow), grass-shrubland, and regenerating native forests (Figure 1). Trophic rewilding over more than 15 years saw the gradual removal of livestock, fencing, hunting, and poaching to restore the puma-guanaco trophic cascade (see Supplementary Information Section S.1 for more details).

**Figure 1.**
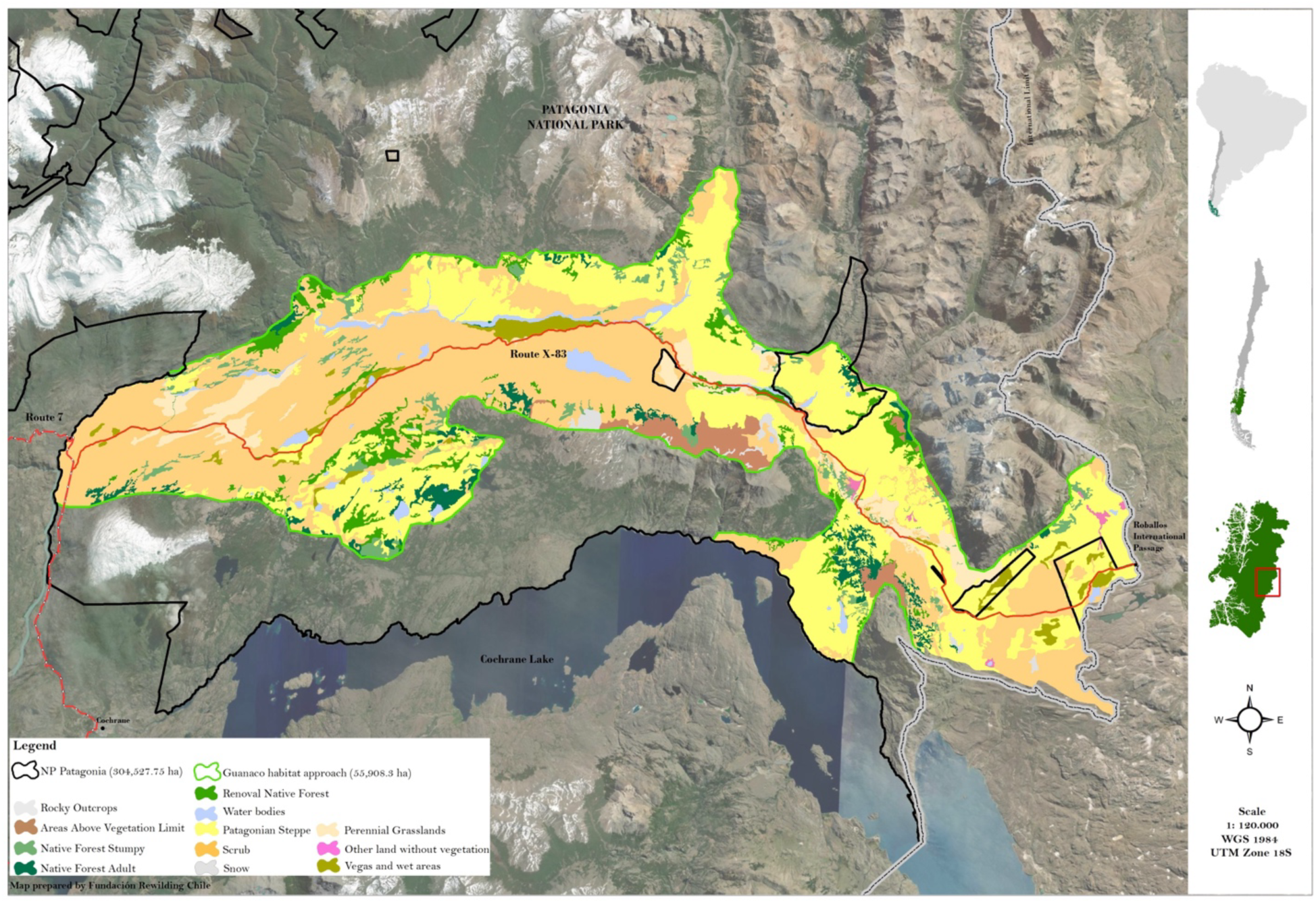
Landcover types within the Valle Chacabuco rewilding area in Patagonia NP, Chile. The landscape includes four grassland types—coironales (Patagonian steppe), grass-shrubland, perennial grassland, and vegas (wet meadow)—juxtaposed with old and young forest, outlined by the guanaco range distribution. The carbon modelling considered the four grassland types in which guanaco feed. These habitat types together cover a spatial extent of 485.91 km^2^.

In principle (Figure 2), by removing plant biomass through herbivory (Donadio et al., 2022), guanacos could reduce the grassland’s capacity to capture and store carbon. However, guanacos can have counteracting positive effects by redistributing and recycling nutrients as feces, urine, and carcasses (Donadio et al., 2022; Bosco et al., 2025). These nutrient inputs can enhance plant production because they are more readily decomposed than plant litter (Holdo et al., 2007). Moreover, guanaco trampling can enhance soil carbon retention (Bosco et al. 2025). Puma predation can control guanaco abundance and foraging, reducing guanaco negative effects of plant biomass (Donadio et al., 2022; LaBarge et al., 2022). Pumas contribute to recycling through their own body waste and prey carcass deposition, potentially offsetting reduced guanaco-mediated nutrient recycling (LaBarge et al., 2022). Field research to empirically resolve how these direct and feedback effects of puma and guanaco impact ecosystem carbon cycling (Figure 2) requires challenging, long-term exclusion experiments. Our modeling experiment thus offers a way to begin resolving each species’ impacts through systematic analysis of three scenarios: (i) an ecosystem with soil and plants only; (ii) an ecosystem with soil, plants, and herbivore (guanaco), and (iii) an ecosystem with soil, plants, herbivores, and predators (puma).

**Figure 2.**
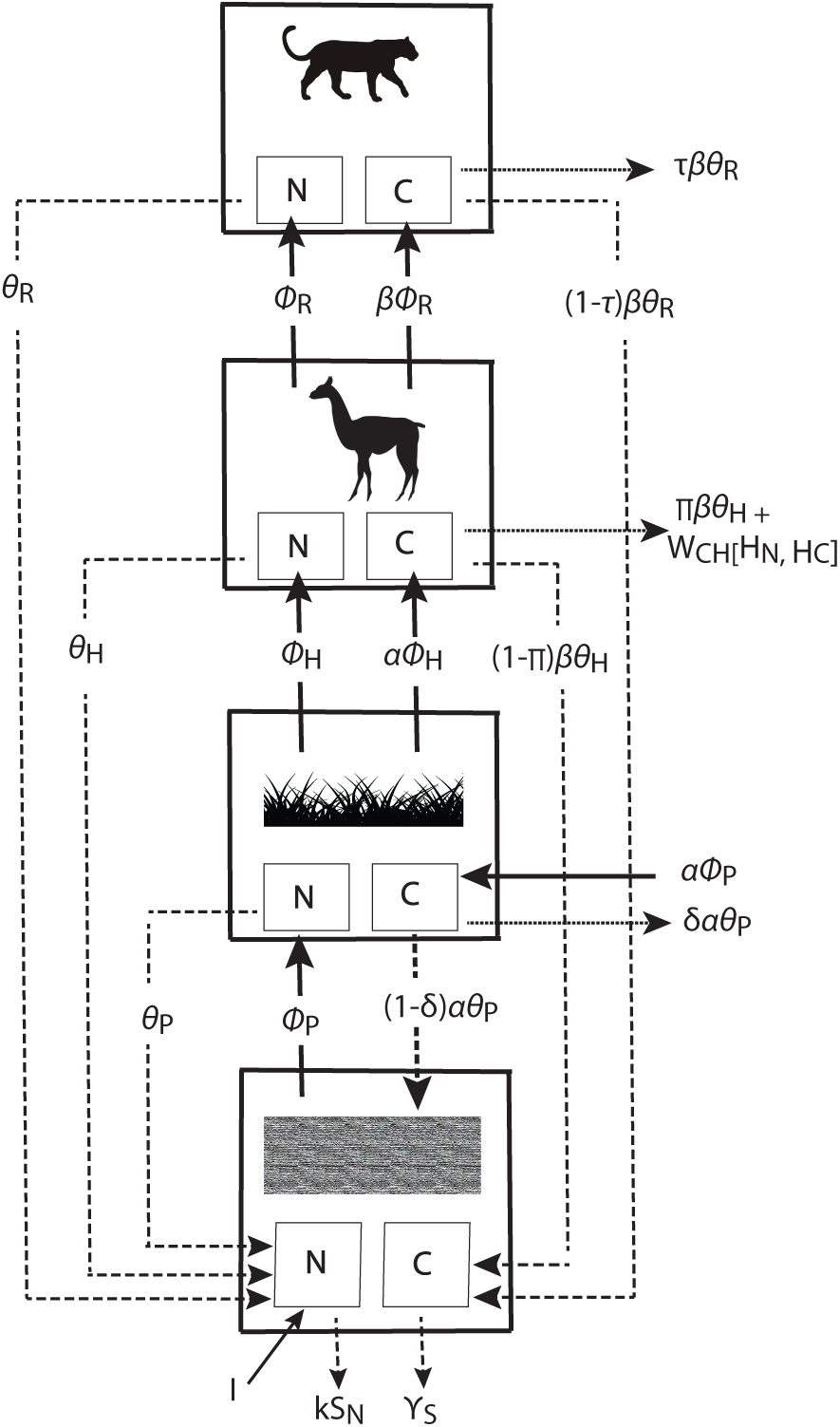
The carbon cycle model applied to the Patagonian grassland ecosystem. The model tracks the flow of carbon (C) and nitrogen (N) between, and N and C stocks in, soil, plant, herbivore (guanaco) and carnivore (puma). Solid arrows depict nutrient uptake by a trophic compartment, dashed arrows depict nutrient recycling, and dotted arrows depict respiratory loss. The model accounts for atmospheric deposition of N to the ecosystem and soil N leaching out of the ecosystem. The modelling further accounts for atmospheric C uptake by grassland plants, plant uptake of soil N, plant C and N by herbivores, herbivore C and N uptake by carnivores. The model accounts for N and C inputs back to soil via litterfall and animal waste and soil, plant, and animal C loss via respiration. Parameters are defined in Tables 1–2.

**Table 1.**
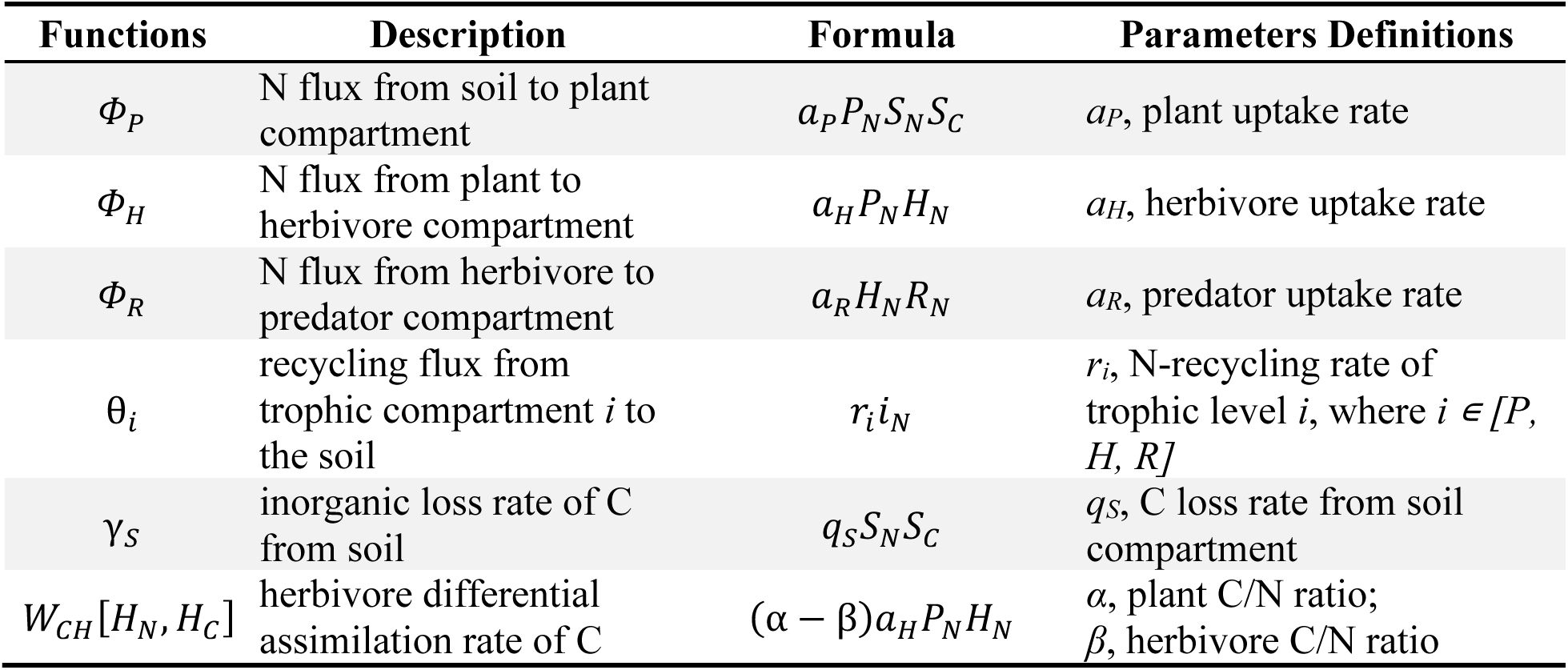
Summary of expressions for plant, herbivore, and predator growth and recycling functions in Equations (1.*a*)–(1.*h*). See Table 2 for units of measurement, parameter definitions, and literature-sourced values. Abbreviations: *N*, nitrogen; *C*, carbon; *S*, soil; *P*, grassland plants; *H*, herbivores (guanaco), *R*, carnivores (puma). See Rizzuto et al. (2024) for the derivation of the *W_CH_*[*H_N_, H_C_*] term.

**Table 2.** Parameter values sourced from the literature used in the final model. See text for scenarios descriptions. See Table S.11 in the Supplementary Information for literature sources for each parameter and information on the adjustments needed to reach a balanced carbon budget and realistic animal density estimates (see section 2.2.1 Model Analysis).

| Description | Scenario | Value | Units | Model parameter | Notes |
| --- | --- | --- | --- | --- | --- |
| Inorganic N inputs | i, ii, iii | $9.30 \times 10^{-7}$ | $\text{kg N m}^{-2} \text{d}^{-1}$ | $I$ | * |
| Plant uptake rate | i, ii, iii | $2.50 \times 10^{-1}$ | $\text{d}^{-1}$ | $a_P$ | ** |
| Herbivore uptake rate | ii | $1.33 \times 10^{-1}$ | $\text{kg N}^{-1} \text{d}^{-1}$ | $a_H$ | *** |
| Herbivore uptake rate | iii | $1.07 \times 10^{-1}$ | $\text{kg N}^{-1} \text{d}^{-1}$ | $a_H$ | *** |
| Predator uptake rate | iii | $8.97 \times 10^{-2}$ | $\text{kg N}^{-1} \text{d}^{-1}$ | $a_R$ | ** |
| Plant respiration rate | i | $1.22 \times 10^{-3}$ | $\text{d}^{-1}$ | *Contributes to $\delta$* | ** |
| Plant respiration rate | ii, iii | $1.09 \times 10^{-3}$ | $\text{d}^{-1}$ | *Contributes to $\delta$* | ** |
| Herbivore respiration rate | ii, iii | $2.42 \times 10^{-2}$ | $\text{d}^{-1}$ | *Contributes to $\pi$* | *** |
| Predator respiration rate | iii | $8.58 \times 10^{-3}$ | $\text{d}^{-1}$ | *Contributes to $\tau$* | *** |
| Plant recycling rate | i | $4.97 \times 10^{-3}$ | $\text{d}^{-1}$ | $r_P$ | ** |
| Plant recycling rate | ii, iii | $3.77 \times 10^{-3}$ | $\text{d}^{-1}$ | $r_P$ | *** |
| Herbivore recycling rate | ii | $5.34 \times 10^{-4}$ | $\text{d}^{-1}$ | $r_H$ | *** |
| Herbivore recycling rate | iii | $1.15 \times 10^{-3}$ | $\text{d}^{-1}$ | $r_H$ | *** |
| Predator recycling rate | iii | $4.30 \times 10^{-5}$ | $\text{d}^{-1}$ | $r_R$ | *** |
| Plant C:N | i, ii, iii | 27.0 | unitless | $\alpha$ | * |
| Animal C:N | ii, iii | 3.29 | unitless | $\beta$ | *** |
| Soil C leaching rate | i | $2.59 \times 10^{-2}$ | $\text{kg N m}^{-2} \text{d}^{-1}$ | $q_S$ | ** |
| Soil C leaching rate | ii | $2.08 \times 10^{-2}$ | $\text{kg N m}^{-2} \text{d}^{-1}$ | $q_S$ | ** |
| Soil C leaching rate | iii | $5.57 \times 10^{-2}$ | $\text{kg N m}^{-2} \text{d}^{-1}$ | $q_S$ | ** |
| Soil N leaching rate | i, ii, iii | $3.50 \times 10^{-4}$ | $\text{d}^{-1}$ | $k$ | ** |

\* Data from ecosystem types in Chile; \*\* Data from related ecosystems in southern South America;
\*\*\* Data from the general literature

The signature for a classic trophic cascade would be lower plant carbon capture and storage in scenario (ii) relative to scenario (i); and higher levels of capture and storage in scenario (iii) relative to scenario (ii). Thus, comparison of the model’s estimates of plant carbon capture and storage between these three scenarios enables resolving how much carbon could be gained or lost by rewilding just guanaco, or guanaco and puma together to this ecosystem.

### 2.2 The carbon cycle model

Previous modelling of predator cascading effects on ecosystem carbon dynamics combined models of wildlife population and woodland vegetation dynamics, and implicitly estimated carbon dynamics by converting plant biomass estimates into plant carbon estimates (Spracklen et al. 2025)—without accounting for animal-driven recycling feedbacks to the soil. We instead modelled carbon dynamics explicitly by using an ecosystem model that includes soil (*S*), plant (*P*), herbivore (*H,* guanaco), and predator (*R,* puma) trophic compartments (Figure 2; Rizzuto et al. 2024). The model is stoichiometrically explicit and uses the carbon to nitrogen ratio (henceforth, C:N) to quantify these nutrients’ stocks (i.e., storage) and fluxes across all trophic compartments (Rizzuto et al., 2024; Schmitz & Leroux, 2020). Equations (1.*a*)–(1.*h*) describe the model ecosystem (Rizzuto et al., 2024):

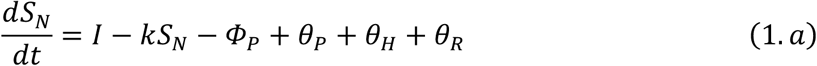

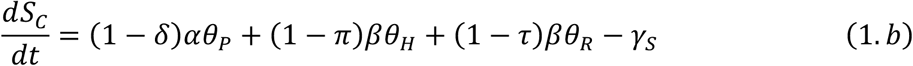

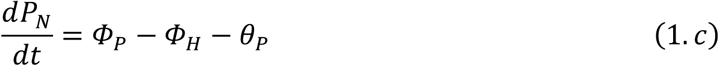

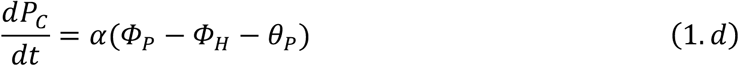

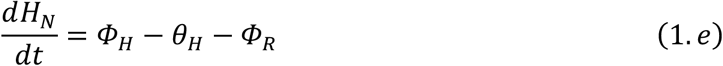

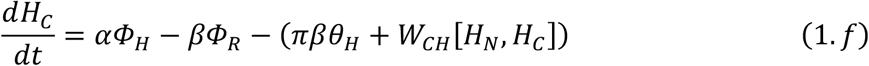

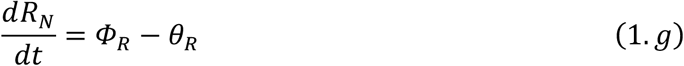

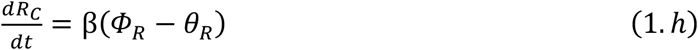

Terms *Φ_i_*and *θ_i_* (*i* ∈ [*S*, *P*, *H*, *R*]), *γ_S_*, and W_CH_[H_N_, H_C_] are described in Table 1. The term W_CH_[H_N_, H_C_] accounts for carbon uptake by herbivores and partitions it between respiration and recycling to the soil depending on the balance of plant and herbivore C:N (Rizzuto et al., 2024). Table 2 presents the model’s parameters, units of measurement, and literature-sourced values. We refer interested readers to Rizzuto et al. (2024) for the model’s structure, dynamics, and numerical analyses.

Equations (1.*a*) – (1.*h*) represent scenario (iii), an ecosystem with a complete food web. Setting parameters *a_R_* and *r_R_* = 0 (Table 2) in *Φ*_R_ and *θ_R_* (Table 1) recovers scenario (ii). Additionally setting *a_H_* and *r_H_* = 0 (Table 2) in *Φ_H_*, *θ_H_*(Table 2) recovers scenario (i). We solved Equations (1.*a*) – (1.*h*) for equilibrium in Mathematica (v. 13.2, Wolfram Research, 2022). The model equilibria are presented in SI Section S.2.

#### 2.2.1 Model Analysis

We ran all analyses in R (v. 4.5.1; R Core Team, 2025). For each scenario, we substituted empirical estimates of parameters (Table 2) into the equilibrium solution to assess how changing the ecosystem’s trophic structure influences plant and soil C stocks (i.e., C storage), net primary productivity (NPP; net C capture by plants), and net ecosystem carbon balance (NECB; ecosystem carbon sink strength) (Rizzuto et al. 2024). We converted steady state C stock estimates for puma and guanaco into whole-animal densities (individuals km^-2^) by dividing carbon stocks by the average adult body mass of puma (61.65 kg; Maher & Moore, 1992) and guanaco (100 kg; Donadio et al., 2020) and assuming animal body mass contains 51% carbon (Franke & Weniger, 1958).

We assumed guanacos spend most of their time foraging in suitable habitat— coironales, grass-shrubland, perennial grasslands, vega (Figure 1)—removing plant biomass C and returning C as body waste (faeces, urine, and body carcasses). A full suite of empirical data values to parameterize the model specifically for each of the habitat types in the trophic rewilding area is yet not available. Therefore, we used empirical measurements from published studies for other Patagonian or Andean grasslands, or similar Chilean ecosystems (Table 2). Thus, our values represent average values for the four habitat types. We parameterized guanaco and puma model components, using data from studies in southern Chile whenever available, and data for these species from general sources including published allometric regression models, when unavailable for the focal system (Table 2). We expressed all parameter values in units of kg, m^2^, and d, reflecting the scales of plant and ecosystem measurements in the literature (Table 2).

Some parameter values are shared across the three modelling scenarios (e.g., inorganic N input rate, soil N leaching rate; Table 2), whereas others change among scenarios—e.g., the values for plant respiration and recycling rates were altered to account for animal impacts (Table 2). We assumed plant recycling rate under herbivory to be 75% of plant recycling rate without herbivory based on measures of plant compensatory changes in litter quality and production under herbivory (i.e., increased leaf lignin and phenolic content; Carrera et al., 2008), and changes in soil physicochemical properties that impact decomposition through trampling (Prieto et al., 2011). We assumed plant respiration rate under herbivory to be 89% of plant respiration rate without herbivory, given evidence from prairie grasslands experiencing bison (*Bison bison*) grazing (Risch & Frank, 2006). We modeled respiration loss for plants and herbivores as proportions of C lost from either trophic compartment through respiration (*δ, π, τ*; Table 2), with the remaining part (i.e., *1-δ*, *1-π, 1-τ*) recycled in the soil compartment. Before running each scenario, we converted raw respiration loss data for compartment *i* ∈ [*P*, *H*, *R*] into proportions as,

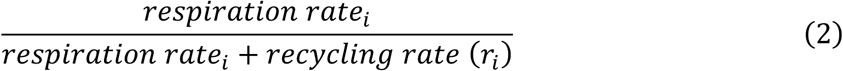

Scenario (i) served as a baseline, simulating animal absence from the ecosystem. To run scenario (ii), an ecosystem with guanaco, we assigned values (Table 2) to herbivore-related parameters that were set to 0 in scenario (i) and updated plant recycling rate (*r_P_*) and carbon loss via plant respiration (*δ*) values, as these plant functional traits change under herbivory (see above). For scenario (iii), we assigned values (Table 2) to predator-related parameters that were set to 0 in scenarios (i) and (ii) and we updated the values of herbivore uptake (*a_H_*) and recycling (*r_H_*) rates as these parameters change due to herbivore vigilance from predation risk (Donadio et al., 2022). See our online repository for code and data used to model all three scenarios.

The equations used to estimate NPP and NECB (kg C m^-2^ d^-1^) are presented in the Supplementary Information (Section S.3 and Table S.1). Consistent with methods used in syntheses of carbon fluxes in ecosystems (Bai & Cotrufo 2022), we scaled daily estimates to yearly estimates assuming a 120-day growing season length for primary productivity, following empirical evidence that soil and plant fluxes are very low to dormant in grasslands outside the growing season (Paruelo et al. 2000). In estimating yearly NECB values, we accounted for enteric methane (CH_4_) emissions from guanaco. No empirical measurement of guanaco CH_4_ emission exist. We thus used allometric regression to estimate annual CH_4_ emission for an average 100 kg adult guanaco (Smith et al., 2015). For yearly estimates encompassing both the growing and non-growing months, we accounted for guanaco and puma CO_2_ respiration for an assumed non-growing season of 245 days.

Parameter values in Table 2 establish an order of magnitude as a basis for modelling. Many parameter values were based on data collected from outside of the focal region, or in some cases measured under laboratory conditions (e.g., plant N uptake and respiration) or estimated from allometric equations (i.e., puma and guanaco body waste release). Because of this, their values had to be adjusted to reach a balanced carbon budget while producing a realistic, feasible solution that approximated guanaco and puma densities observed for Patagonian grasslands (1.1–30.2 guanacos/km^2^ throughout Patagonia, (Carmanchahi et al. 2022); 0.034–0.075 pumas/km^2^, (Reinhart et al., 2014)). Six parameters required adjustment, including soil leaching rates of carbon (*q_S_*) and nitrogen (*k*), guanaco (*a_H_*) and puma (*a_R_*) nutrient uptake rates, plant litter recycling rate (*r_P_*), plant respiration rate, and guanaco (*r_H_*) and puma (*r_R_*) recycling rates. This practice is consistent with other modelling of animal impacts on ecosystem biogeochemistry (e.g., Holdo et al., 2007).

#### 2.2.2 Model Sensitivity and Uncertainty Analysis

We conducted sensitivity analyses to examine how model estimates of NPP, NECB, and soil and plant C stocks varied with changes in the values of the six key parameters that were adjusted. Of these, *q_S_*, *r_P_*, and plant respiration rate appear in all modelling scenarios, *a_H_* and *r_H_* appear in scenarios (ii) and (iii), and *a_R_* and *r_R_* appear only in scenario (iii). We analysed the model’s sensitivity to each parameter by varying the focal parameter by ±30% the value used in the model while keeping all other parameters fixed at their literature-sourced value. For each parameter, we randomly sampled 10000 values from this range and then ran the model scenarios in which the focal parameter appeared.

We used these sensitivity analyses to estimate uncertainty around model estimates of NPP, NECB, and soil and plant C stocks in the presence of the animals. From the broader sensitivity analyses, we extracted those estimates that produced guanaco and puma equilibrium density estimates within the ranges observed in the field, i.e., 1.1–30.2 guanacos/km^2^ (Carmanchahi et al. 2022) and 0.034–0.075 pumas/km^2^ (Reinhart et al., 2014). We then characterized uncertainty as a range determined by 2× the standard deviation of the estimates of NPP, NECB, and soil and plant C obtained for the range of equilibrium animal densities. See SI Section S.6 and the Supplementary Code document in our repository for details.

## 3 RESULTS

At steady state (balanced carbon budget), the mean carbon capture and storage estimates were associated with a density of 20.3 guanaco/km^2^ in scenario (ii), and 30.9 guanaco/km^2^ in the presence of 0.0124 pumas/km^2^ for scenario (iii) (Table S.2).

### 3.1 Estimated carbon capture

The modelling revealed trophic cascading effects on ecosystem carbon capture (Figure 3, Table S.4). On average, estimated NPP decreased by 38% between scenarios (i) and (ii). NPP increased by 2.69× between scenarios (ii) and (iii). Moreover, the model estimated that NPP could be 1.68× higher in scenario (iii) than in scenario (i). NECB (Figure 3), albeit lower than NPP in scenarios (ii) and (iii), shows the same qualitative trends as NPP.

**Figure 3.**
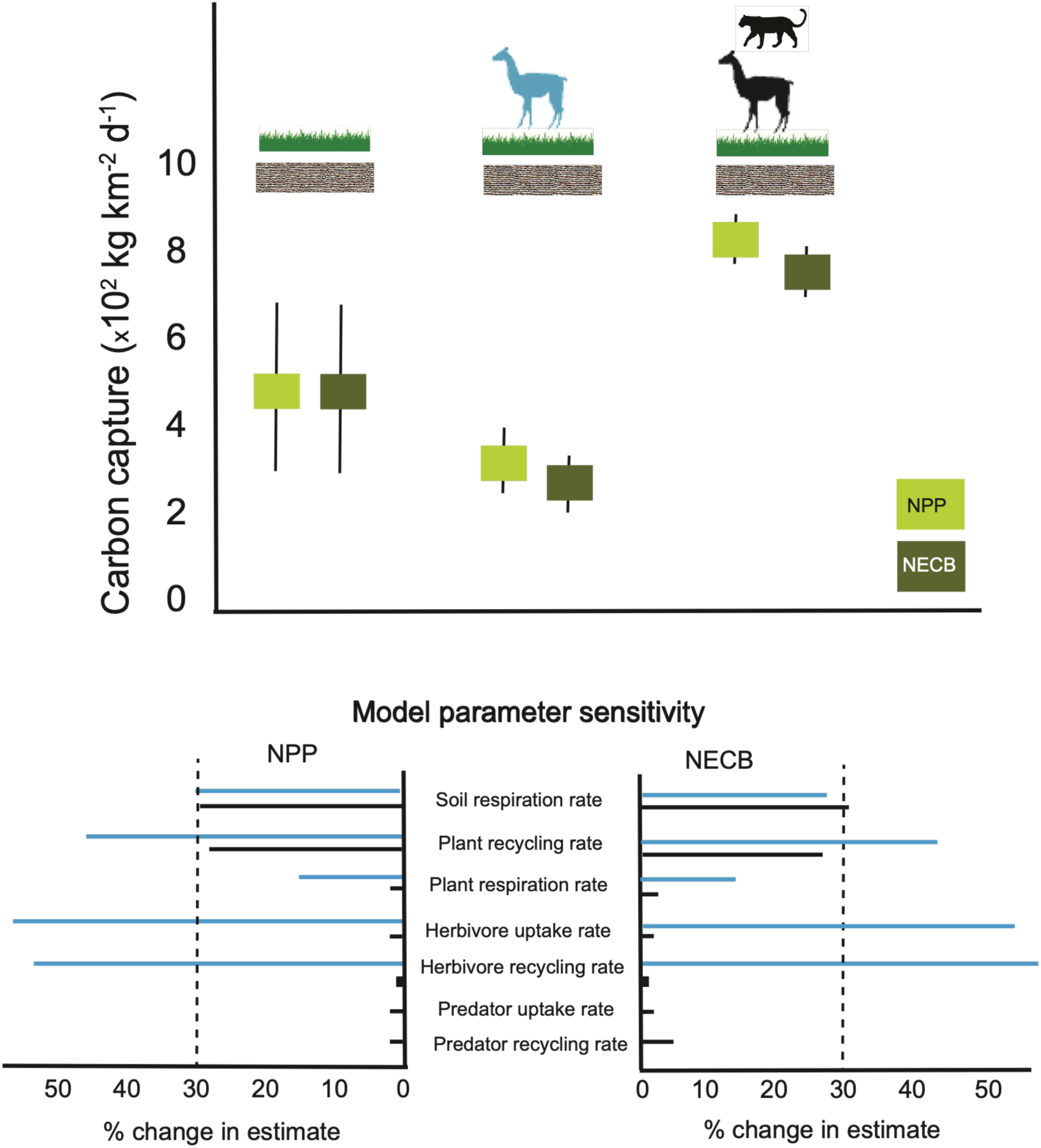
Model estimated means and uncertainty ranges (± 2 × SD of model estimates) for Net Primary Productivity (NPP) and Net Ecosystem Carbon Balance (NECB) among scenarios (i), (ii), and (iii). Model estimates for soil and plant carbon storage are variously sensitive to 30% change in each model parameter value. Blue lines are sensitivities for scenario (ii) and black lines are for scenario (iii). Percentage changes in model estimates exceeding 30% (vertical dashed line) are considered highly sensitive; values less than 30% are less sensitive or insensitive (% change in estimates less than 2%).

The model estimates can vary considerably due to model sensitivity to parameter changes (Figure 3, Tables S.5–S.10). In scenario (ii), the guanaco’s estimated effects on NPP and NECB were most sensitive (i.e., the % change in the estimate was ≥ 30%) to changes in soil respiration rate, plant recycling rate, and guanaco nutrient uptake and recycling rates (Figure 3). In scenario (iii), the estimated effects of guanaco and puma together on NPP and NECB were sensitive to changes in soil respiration and plant recycling rates, but less so or invariant to changes in other parameters (Figure 3).

The sensitivity to changes in model parameters led to uncertainty in model estimates of carbon capture (Figure 3). Uncertainty was greatest in scenario (i) and diminished with increasing numbers of trophic levels in the system (Figure 3). The uncertainty range for scenario (i) overlapped with the estimated mean and uncertainty range for NPP in scenario (ii), suggesting there are conditions where NPP may not decrease in the presence of guanacos. However, the lower estimates for NECB in scenario (ii) did not overlap with scenario (i) because scenario (ii) accounts for additional carbon losses from animal respiration and CH_4_ release. Uncertainty in NPP and NECB estimates for scenario (iii) were quite low and did not overlap with the other scenarios. See SI Section S.6 and Tables S.5–S.10 for median estimates and ranges of ecosystem carbon capture from the sensitivity analyses.

### 3.2 Estimated carbon storage

Consistent with expectations for a trophic cascade, the estimated plant carbon stock decreased on average by 15% between scenarios (i) and (ii), but increased by 2.63× between scenarios (ii) and (iii) (Table S.3). With both animals present, the model estimated that the ecosystem could store 2.23× more C in plant biomass than in their absence. The model estimates of plant carbon stocks were sensitive to guanaco nutrient uptake and recycling rates in scenario (ii). The estimated effects of puma and guanaco in scenario (iii) were most sensitive to soil respiration rate and less so for all other parameters.

The estimated average soil carbon stock decreased by 10% between scenarios (i) and (ii), but it increased slightly by 1.02× between scenarios (ii) and (iii) (Table S.3). Overall, the model estimated that the ecosystem could have 25% less soil C storage with than without both animals. Guanaco’s estimated effects on soil C in scenario (ii) were most sensitive to changes in soil respiration rate, plant recycling rate, and guanaco nutrient uptake and recycling rates. The estimated effects of guanaco and puma together (scenario (iii)) were sensitive to plant recycling rate and less so or invariant to changes in other parameters.

When scaled as a percentage of the mean estimate, the magnitude of uncertainty in model estimates of soil carbon stocks was 1.3× larger than for plant carbon stocks (Figure 4). Uncertainty again decreased with increasing numbers of trophic levels in the ecosystem. There was no overlap in the uncertainty range for estimated plant carbon stocks among scenarios. For soil carbon stocks, the uncertainty range for scenario (i) overlapped with the estimated mean and uncertainty range in scenarios (ii) and (iii), suggesting there could be conditions where there may be no differences in soil C stocks between animal absent vs present scenarios.

**Figure 4.**
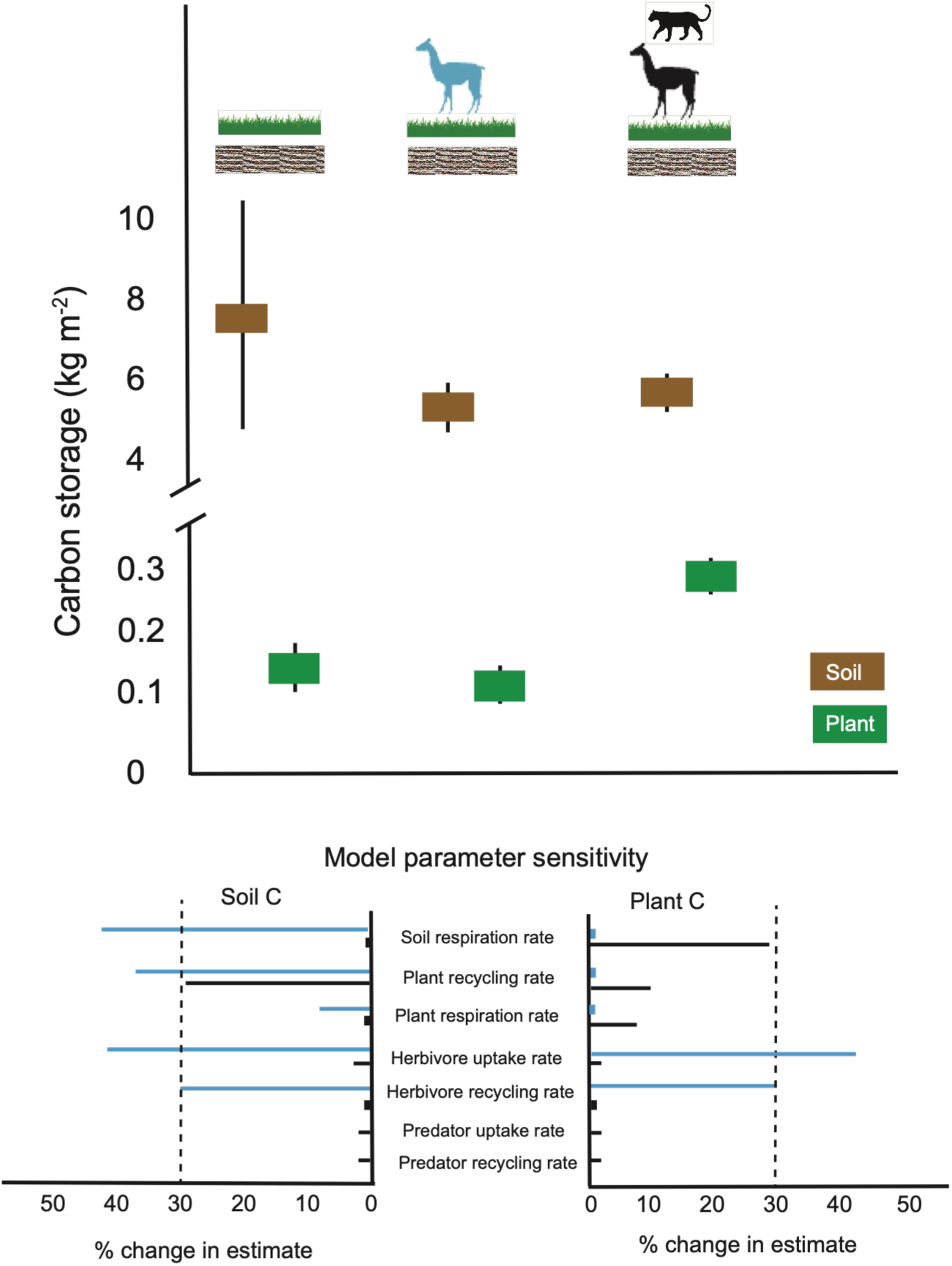
Model estimated means and uncertainty ranges (± 2 × SD of model estimates) for soil and plant carbon storage among scenarios (i), (ii), and (iii). Model estimates for soil and plant carbon storage are variously sensitive to 30% change in each model parameter value. All specifications as in Figure 3.

### 3.3 Validating carbon capture and storage estimates

Measurements of puma- and guanaco-driven carbon capture and stocks do not exist yet, so estimates for scenarios (ii) and (iii) cannot yet be validated. However, the model can be validated by comparing estimates for scenario (i) against field measures of NPP and carbon stocks for Patagonian grasslands. NPP measurements reveal that Patagonian grasslands can capture 7.72 × 10^1^–1.03 × 10^3^ kg C km^-2^ d^-1^ (Peri et al. 2015), a range containing the model’s uncertainty range in estimated NPP of 3.38 × 10^2^–6.94 × 10^2^ kg C km^-2^ d^-1^ (Figure 3).

Consistent with published field measurements for southern Chilean and Patagonian grasslands (Perez-Quezada et al., 2022; Peri et al., 2018), the model estimated that at steady state plant carbon stocks should be ∼1 order of magnitude less than soil carbon stocks (Figure 4). The uncertainty range of model estimated plant carbon stock (0.09–0.17 kg C m^-2^; Figure 4) is conservative, given a measured value of 0.353 kg C m^-2^ in plant biomass in southern Chilean grassland (Perez-Quezada et al. 2022). The uncertainty range of 4.9–10.1 kg C m^-2^ in estimated soil carbon stock (Figure 4) aligns with the 4.54–11.24 kg C m^-2^ range measured in shrub-steppe, grass-steppe, and grassland habitats in southern Patagonia (Peri et al. 2018).

## 4 DISCUSSION

Trophic rewilding is being partly promoted to rebuild ecosystem trophic structure and reinstate trophic cascades to restore the vegetation structure of ecosystems (Svenning et al., 2019; 2024). Yet, the consequences of restoring trophic cascades beyond plants on whole ecosystem biogeochemical and carbon cycling are often unexplored and thus remain poorly understood (Schmitz et al., 2018). Developing understanding requires a holistic conceptualization of ecosystems that—alongside the consumptive relations among predators, herbivores, and plants (Terborgh et al., 2010)—includes soil trophic compartments and animal recycling feedbacks to them (Schmitz et al., 2018; Schmitz & Leroux, 2020).

Our holistic modelling revealed that restoring a trophic cascade could indeed propagate changes in carbon capture and storage in plants, consistent trophic cascade theory (Figures 3 and 4). The modeling revealed that an intact Patagonian grassland ecosystem that includes pumas and guanacos could increase carbon capture (NPP and NECB) by 1.27×– 2.5× above that for the ecosystem without the animals. Plant carbon storage is estimated to be 1.76×–3.25× higher in the intact ecosystem than in the absence of animals. These increases compare with modelling estimates of trophic cascade effects of wolves (Spracklen et al. 2025) and empirical measures for tigers (Roberts et al., 2025). However, soil carbon dynamics are more complex, and the intact ecosystem could enhance soil carbon storage by up to 1.16× or reduce it by up to 0.57×.

On an annual basis, which accounts for carbon gains during a 120 d growing season and carbon losses from CO_2_ respiration and enteric methane releases in the 245 d nongrowing season, the modelling suggests that NECB could amount to 94,405 kg C per km^2^ (uncertainty range 94,375 kg C km^-2^–94,435 kg C km^-2^) of which 23%–43% attributable to animal effects. In the intact ecosystem, steady state plant biomass storage could reach 2.9 × 10^5^ kg C km^-2^ (2.8 × 10^5^ kg C km^-2^–3.0 × 10^5^ kg C km^-2^) of which 43%—67% attributable to animal effects. Similarly, estimated steady state soil storage with animals could amount to 5.7 × 10^6^ kg C km^-2^ (5.6 × 10^6^ kg C km^-2^–5.8 × 10^6^ kg C km^-2^)—i.e., an average loss of 1.78 × 10^6^ kg C km^-2^ relative to the no-animal scenario. However, given the wide uncertainty of soil C stock estimates in the absence of animals, the intact ecosystem’s soil stock could gain 1.78 × 10^5^ kg C km^-2^; or lose up to 4.3 × 10^6^ kg C km^-2^. Empirical analyses focused on soil processes with and without animals are necessary to obtain a reliable, whole-ecosystem carbon budget. The modelling also showed that adding trophic levels to the modelled ecosystem reduced variability around the average model estimates, implying that animals may contribute to stabilizing the amount of carbon captured and stored in plants and soils in ecosystems (Naidu et al., 2022).

Counterintuitively, the model predicts more carbon captured and stored in plant biomass on average when both species are present than when they are absent. Under a balanced carbon budget, the modelling also predicts 30.9 guanaco/km^2^ with pumas on average—1.5× higher than average guanaco density without pumas (20.3 individuals/km^2^). Yet, both outcomes are plausible, being observed in experiments that traced predator presence’s effects on ecosystem productivity and prey abundance (Strickland et al., 2013). In our model, predators’ effects are explained by changes in the pathway followed by nutrients in plant biomass. Instead of being completely recycled as plant litter, some standing plant biomass is diverted via consumption and trophic transfer among the animal trophic compartments and then recycled as animal body waste (Figure 2). This animal-driven recycling feedback can increase plant productivity relative to that supported by plant-litter-derived nutrients (Schmitz et al., 2018). However, guanaco presence reduces plant productivity and carbon storage, consistent with guanaco consumptive effects on plants outweighing the recycling effects on plant productivity. Adding pumas to the model ecosystem reverses this effect of guanaco, and nutrient recycling from pumas and guanacos together enhances plant productivity beyond the no-animal scenario. Hence, the modelled pumas are “engineering” their ability to be part of the ecosystem via their added nutrient cycling feedback effects that enhance primary productivity of this comparatively unproductive system and lead to high-enough guanaco densities to support the puma population. This result is consistent with model evidence that steady state marine fish prey abundances were higher with top predators than without, due to higher productivity supporting both predators and prey (Christensen & Pauly, 1998). It also agrees with classic theoretical predictions that, to support predators and prey together, ecosystems must reach higher productivity levels than those needed to support prey alone (Oksanen et al., 1981).

The modelling revealed that restoring a trophic cascade did not carry to soil storage, with an estimated average 24% less carbon stored in soils with vs without animals (Figure 4). Even accounting for plant carbon gains due to puma and guanaco presence results in an estimated average 20% less carbon storage per unit area when animals are present. This reinforces recent empirical findings that considering animal effects merely on aboveground plant carbon can give inaccurate insights about the net effects of animals on whole ecosystem carbon storage (e.g., Kindermann, et al. 2025; Roberts et al, 2025).

Conclusions about trophic cascade impacts based on the estimated average responses across trophic levels need to be drawn cautiously. Model estimates are highly sensitive to changes in several parameters values, especially soil respiration, plant recycling, and herbivore nutrient uptake and recycling rates (Figures 3–4). Thus, there may be conditions under which no differences emerge in carbon capture and storage between animals present vs. absent, notably for scenario (i) vs (ii). Some of the uncertainty in model estimates arises because puma’s and guanaco’s effects on some biogeochemical processes have not been measured. For instance, while measures of herbivore consumption rates in relation to vegetation quality exist (Meyer et al. 2010), few studies quantify herbivore effects on plant recycling as it requires measuring plant litter entering the soil with and without animals (Lienau et al. 2026).

The modelling reveals that soil is a key reservoir for carbon in grassland ecosystems. However, our model treats soil as a “black box”—modeling carbon and nitrogen input and output to soil but abstracting within-soil dynamics (Figure 2). Generally, the magnitudes of animal effects on soil processes—due to herbivory, organic matter deposition of body wastes (faeces, urine, peri-natal fluids, carcasses, and body parts), trampling, and bioturbation are poorly characterized and measured, and thus remain highly uncertain (Lienau et al. 2026). Future analyses would benefit from explicitly combining above- and below-ground processes, for example, by leveraging modelling principles used in analyses of belowground biogeochemical cycling (Lienau et al. 2026), to mechanistically elaborate processes currently abstracted in our model. This includes accounting for different forms of dead organic biomass in plant litterfall (e.g., structural vs metabolic litter), uptake and decomposition of animal and plant organic matter by different functional types of soil microbes, and release of elements in organic and inorganic form (Lienau et al. 2026). Thus, more empirical research focused on measuring animal influences on below-ground processes will be critical to fully understand wildlife effects on whole ecosystem carbon dynamics.

The rewilding program in the PNP landscape is restoring grassland ecosystems from livestock to natural grazing by removing 25,000 sheep (52 sheep/km^2^) and 3,000 cattle (6.24 cattle/km^2^) to enable the recovery of guanacos, among other wildlife species. Between 2005–2022, the guanaco population grew rapidly from 900 to 3,000 individuals (Fundación Rewilding Chile, unpublished data). However, sheep and guanacos are often considered functionally equivalent in their grazing impacts (Fernández et al., 2021). Hence, both species have negative impacts on ecosystem carbon capture and storage relative to their absence, beyond densities of 12–14 sheep/km^2^ and ∼9.9 guanacos/km^2^ (Oliva et al. 2019; Peri et al. 2016). Thus, using sheep ranching as an NbCS would require managing animals at such low densities that herding likely would become economically unviable. Our modelling suggests positive carbon benefits could be obtained by enlisting natural processes in a fully intact ecosystem with guanacos and pumas. Such natural processes would sustain higher guanaco densities (30.9 guanacos/km^2^), and thereby the joint recycling effects of pumas and guanacos, which enhance grassland productivity and carbon capture and storage. This would align with the conservation ethic of trophic rewilding, to minimize human management and let nature take its course (Svenning et al., 2024).

We show how trophic rewilding could promote wildlife recovery and conservation in local ecosystems to mitigate climate change and biodiversity loss together. Solutions to each have historically been treated independently, yet this thinking is changing (Schmitz et al., 2023; Svenning et al., 2019; 2024). Hence, protecting animals while ignoring their potential impacts on carbon capture and storage could risk over- or under-estimating the amount of carbon that can be captured and stored in ecosystems. In turn, this risks creating inaccurate carbon budgets, which is problematic given growing interest to develop payment schemes to support local stewardship and help mitigate climate warming by rewilding animals and their ecosystems. The timeline for achieving these outcomes remains generally unknown due to the need for long-term measurement (but see SI Section S.5; Kaštovská et al., 2024; Naidu et al. 2022). However, a priori modelling showing the feasibility of restoring a trophic cascade as an NbCS can help to identify when conducting such long-term measurement efforts would be worthwhile.

In conclusion, we have used a new, animated carbon cycle model to estimate animal effects on ecosystem carbon budgets and provide first approximations of the feasibility of using specific, on-the-ground trophic rewilding projects to enhance carbon capture and storage. The model offers a crucial foundation to assist conservation decision-making aimed at rewilding nature to mitigate climate change and reverse biodiversity loss. Our analyses revealed that rewilding guanaco and pumas together could enhance net ecosystem carbon storage far above what would be sequestered in their absence. Of course, the actual performance of the rewilded ecosystem to capture and store carbon will have to be monitored and verified in PNP using exclosure studies that compare ecosystem responses with and without animals. Regardless, our results offer insights to consider trophic rewilding as a feasible NbCS in Patagonian grasslands.

## 5 ACKNOWLEDGEMENTS

We are grateful to Guillermo Sapaj for useful comments on earlier versions of this manuscript. We are grateful to Fundación Rewilding Chile for their support.

## 6 AUTHORS’ CONTRIBUTIONS

MR, OJS, and CS conceived the idea and designed the study. MR and OJS assembled the data from the literature to parameterize the model. MR led model analyses, calibration, and sensitivity analyses. MR and OJS interpreted the results and led manuscript writing. All authors reviewed and approved the final manuscript.

## 7 DATA AVAILABILITY STATEMENT

All data and code used in the analyses are available in our online repository at: https://figshare.com/s/5d4870da93c39a7efdf9 [note: link is anonymized for peer review, will be updated upon acceptance]

