## Supplementary Information for "Trophic rewilding pumas and guanacos: estimating the potential to enhance carbon sequestration in a Patagonian grassland ecosystem"

### S.1. The study area

Guanaco was once the dominant ungulate grazer in Valle Chacabuco ecosystem for ~10,000 years (Hernandez et al., 2019), but its abundance diminished across the Valle Chacabuco landscape after humans transformed it for livestock (especially sheep) grazing, because native ungulates were perceived to compete with livestock (Hernandez et al., 2019). The erection of extensive fences further reduced wild ungulate movements (Tompkins 2024). Predator control measures to protect livestock reduced mammalian and avian predator numbers, eroding the original functional trophic structure of the ecosystem (Donadio et al., 2022). The trophic rewilding initiative—partially intended to re-establish the puma-guanaco trophic cascade—spanned more than 15 years and saw gradual removal of all livestock, dismantling of hundreds of kilometres of fencing, prohibition of hunting, and poaching controls. These measures enabled wildlife to passively redistribute across the landscape and grow in population size via natural demographic processes. Whether the re-establishment of the puma-guanaco trophic cascade could impact carbon cycling in the grassland ecosystem remains unknown.

### S.2. The model's equilibrium solutions.

We explore the potential for trophic rewilding guanaco and puma in Patagonia NP using the equilibrium solutions (i.e., a balanced carbon and nitrogen budget ) for each of the three modeling scenarios in Rizzuto et al. (2024): (i) carbon uptake and storage in the absence of animals; (2) carbon uptake and storage in the presence of guanaco; (3) carbon uptake and storage in the presence of guanaco and puma. As we are interested in how the presence of herbivores and predators shapes ecosystem carbon capture in Patagonia NP, we compare results of these three modeling scenarios. The ecosystem model we used includes a food chain comprising four compartments: soil, plants, guanaco, and puma. For each of these compartments, the model tracks gains (e.g., inorganic inputs, nutrient uptake, photosynthesis) and losses (e.g., leaching, respiration, recycling). At the ecosystem level, the model estimates gross and net primary productivity and net ecosystem carbon balance (NECB). NECB captures the net accumulation of carbon in ecosystems (Chapin et al., 2006). In measuring NECB, we use an expanded formula that accounts for the direct and indirect effects of animals on ecosystem carbon cycling (see Appendix A.2 and Rizzuto et al. (2024) for more details).

The three model scenarios presented in Rizzuto et al. (2024) have the following sets of equilibrium solutions.

Scenario (i),

$$Soil_N = \frac{I}{k} \tag{S.1}$$

$$Soil_C = \frac{k r_P}{a_P I} \tag{S.2}$$

$$Plant_N = \frac{q_S}{a_P \alpha (1-\delta)} \tag{S.3}$$

Scenario (ii),

$$Soil_N = \frac{I}{k} \qquad (S.4)$$

$$Soil_C = \frac{kr_H r_P(\alpha(1-\delta) + (\pi - 1)\beta)}{I(a_H q_S + a_P(\pi - 1)\beta r_H)} \qquad (S.5)$$

$$Plant_N = \frac{r_H}{a_H} \qquad (S.6)$$

$$Herbivore_N = -\frac{r_P\left(a_H q_S + a_P r_H \alpha(\delta - 1)\right)}{a_H(a_H q_S + a_P(\pi - 1)\beta r_H)} \qquad (S.7)$$

Scenario (iii),

$$Soil_N = \frac{I}{k} \qquad (S.8)$$

$$Soil_C = \frac{k(a_R r_P + a_H r_R)}{a_P a_R I} \qquad (S.9)$$

$$Plant_N = -\frac{a_R q r_P + a_H q r_R + a_P r_H r_R \beta(\pi - \tau)}{a_P\left(a_R r_P \alpha(-1+\delta) + a_H r_R \beta(-1+\tau)\right)} \qquad (S.10)$$

$$Herbivore_N = \frac{r_R}{a_R} \qquad (S.11)$$

$$Predator_N = -\frac{a_H a_R q r_P + a_H^2 q r_R + a_H a_P(-1+\pi) r_H r_R \beta + a_P a_R r_H r_P \alpha(-1+\delta)}{a_P a_R\left(a_R r_P \alpha(-1+\delta) + a_H r_R \beta(-1+\tau)\right)} \qquad (S.12)$$

Across all three scenarios, N represents nitrogen and C represents carbon. *S*, *P*, *H*, and *R* represent soil, plants, herbivores, and predators, respectively. Parameters are briefly described in Table 1 in the main text, and in more detail in Rizzuto et al. (2024). The model uses C:N (parameters $\alpha$ for plants and $\beta$ for animals) to track the budget and cycling of C in the ecosystem and account for the rate limitation imposed on them by the cycling of an essential nutrient, N. Hence, the C content of plant, herbivores, and predators is recovered as,

$$Plant_C = \alpha \cdot Plant_N \qquad (S.13)$$

and

$$Herbivore_C = \beta \cdot Herbivore_N \qquad (S.14)$$

$$Predator_C = \beta \cdot Predator_N \qquad (S.15)$$

### S.3. Net Ecosystem Carbon Balance formula derivation.

Net Ecosystem Carbon Balance (NECB) measures ecosystem carbon storage in terms of the net difference between an ecosystem's anabolic and catabolic processes, i.e., the balance between net rate of carbon accumulation in ecosystems due to carbon fixation by plants (primary productivity), heterotrophic (i.e., animals, microbes) production and respiration, as well as additional losses including methane emissions directly from animals and soils and sediments of ecosystems (Chapin et al., 2006; Chapin III et al., 2002). Traditionally, net ecosystem carbon balance is estimated as

$$NECB = Gross\ Primary\ Production - Ecosystem\ Respiration - Lateral\ Fluxes \quad (S.16)$$

where Gross Primary Production is essentially gross carbon uptake by plants, Ecosystem Respiration comprises respiration by all trophic compartments (e.g., soil, plants, herbivores, predators; Chapin et al., 2006), and Lateral Fluxes comprise losses of C through mechanisms other than respiration (e.g., leaching to groundwater, methane emissions). However, by only accounting for the effects of heterotrophs through respiration, equation S.16 does not account for the direct and indirect effects of animals on an ecosystem's trophic compartments, including net assimilation of C in animal biomass (secondary productivity that varies with plant and animal stoichiometry) and N and C release from animals (recycling feedbacks that also vary with animal stoichiometry) that promote Gross Primary Production, thereby missing important contributions to an ecosystem's C budget by these actors via the processes they mediate.

Hence, Rizzuto et al. (2024) expanded on equation S.16 to integrate the effects of heterotrophs in the accounting of NECB. Broadly, Rizzuto et al. (2024) defined NECB as the algebraic sum of Net Primary Production (NPP) and Net Heterotrophic Production (NHP),

$$NECB = NPP + NHP \quad (S.17)$$

These two components capture the combined anabolic and catabolic processes happening in an ecosystem, across all trophic compartments. Equation S.17 is conceptually comparable to equation S.16 but, by explicitly considering net heterotrophic production, it allows for debiting each trophic compartment's heterotrophic respiration from its own heterotrophic production. Net Primary Production is the balance of the photosynthetic and respiratory processes that happen in the autotroph compartment of an ecosystem. If we imagine a terrestrial ecosystem, where primary producers are generally plants, we can then specify:

$$NPP = Gross\ Plant\ Production - Plant\ Respiration \quad (S.18)$$

Conversely, Net Heterotrophic Production is the algebraic sum of all biomass-producing and respiratory processes taking place in the heterotrophic compartments of the ecosystem. These include (i) any trophic level above the autotrophs—e.g., herbivores, predators—but also (ii) any trophic level involved in the decomposition pathways that recycle nutrients from waste and dead biomass and make them available to autotrophs once again—the so-called "brown food web". Thus, NHP is (Rizzuto et al., 2024):

$$NHP = (Predator\ Gross\ Production - Predator\ Respiration) + (Herbivore\ Gross\ Production - Herbivore\ Respiration) + (Soil\ Gross\ Production - Soil\ Respiration) \quad (S.19)$$

where Gross Production is measured as the C uptake rate by a given trophic compartment. Accounting for both components of heterotrophic C effects allows us to explicitly measure the relative impact of different kinds of heterotrophs on NECB. In the study, guanaco is the herbivore and puma is the predator.

Table S.1 below reports the formulae used to estimate both NECB and Net Primary Production in our study. Note that all quantities are expressed in units of kg C $km^{-2}$ $d^{-1}$.

Table S.1. Net primary productivity (NPP) and net ecosystem carbon balance (NECB) expressions used in each modeling scenario. Scenarios are: (i) an ecosystem with soil and grassland plant trophic compartments only; (ii) an ecosystem with soil, plant, and herbivore (guanaco), and (iii) an ecosystem with soil, plants, herbivores, and predators (puma). See Table 2 in the main text for parameter definition, values, and units of measurement. All units are kg C km$^{-2}$ d$^{-1}$.

| Scenario | (i) | (ii) | (iii) |
|---|---|---|---|
| NPP | $\frac{q_S r_P}{a_P}$ | $-\frac{r_H r_P \alpha(a_P r_H(\alpha+(-1+\pi)\beta)(-1+\delta)+a_H q_S \delta)}{a_H(a_H q_S + a_P(-1+\pi) r_H \beta)}$ | $\frac{\alpha(-a_H r_R + a_R r_P(-1+\delta))(a_R q_S r_P + a_H q_S r_R + a_P r_H r_R \beta(\pi-\tau))}{a_P a_R(a_R r_P \alpha(-1+\delta) + a_H r_R \beta(-1+\tau))}$ |
| NECB | $\frac{q_S r_P}{a_P}$ | $-\frac{r_H r_P(a_P r_H \alpha^2(-1+\delta)+a_H q_S(\beta-\pi\beta+\alpha\delta))}{a_H(a_H q_S + a_P(-1+\pi) r_H \beta)}$ | $\frac{a_R^2 q_S r_P^2 \alpha(\delta-1) + a_R r_P r_R\left(-a_P r_H \alpha\beta(\delta-1) + a_H q_S(\alpha(\delta-2)+\beta(\tau-2))\right) + a_H r_R^2(a_P r_H \beta(\beta-\pi(\alpha+\beta)+\alpha\tau) - a_H q_S(\alpha+2\beta-\beta\tau))}{a_P a_R(a_R r_P \alpha(\delta-1) + a_H r_R \beta(\tau-1))}$ |

### S.4. Model estimates of animal densities, soil and plant stocks, and ecosystem carbon capture

Here, we present the model's guanaco and puma density (Table S.2), soil and plant carbon stocks (Table S.3), and carbon capture (Table S.4) estimates.

Table S.2. Model estimates of biomass, density, and population size for guanaco and puma for modelling scenarios (ii) and (iii), for the 485.91 $km^2$ Valle Chacabuco ecosystem in Patagonia NP, Chile.

| | **Scenario** | **Biomass** (kg C $m^{-2}$) | **Density** (# ind $km^{-2}$) | **Population size** (# ind)* |
|---|---|---|---|---|
| Guanaco | (ii) | $1.03 \times 10^{-3}$ | 20.3 | 9861 |
| | (iii) | $1.58 \times 10^{-3}$ | 30.9 | 15026 |
| Puma | (iii) | $3.90 \times 10^{-7}$ | $1.24 \times 10^{-2}$ | 6.02 |

*Estimated over a 485.91 $km^2$ subset of total guanaco-suitable habitat in Patagonia NP.

Table S.3. Model Estimates of C stocks in the model's soil and plant compartments, across modelling scenarios for the Valle Chacabuco ecosystem in Patagonia NP, Chile.

| **Scenario** | **Soil** (kg C $m^{-2}$) | **Plant** (kg C $m^{-2}$) |
|---|---|---|
| (i) | 7.5 | 0.13 |
| (ii) | 5.7 | 0.11 |
| (iii) | 5.8 | 0.29 |

Table S.4. Model estimates of net primary productivity (NPP), and net ecosystem carbon balance (NECB, i.e., carbon sequestration) for the Valle Chacabuco ecosystem in Patagonia NP, Chile. The estimates are for the no animal (i), guanaco-only (ii), and guanaco-puma (iii) scenarios. Seasonal and yearly estimates are calculated over an assumed growing season of 120 days. Daily, seasonal, and yearly NECB estimates account for guanaco methane ($CH_4$) emissions, and yearly estimates also account for guanaco and puma respiration for the non-growing part of the year (245 days).

| | **Scenario** | **Growing season** | | **Yearly*** ($kg\ C\ km^{-2}\ y^{-1}$) |
|---|---|---|---|---|
| | | **Daily** ($kg\ C\ km^{-2}\ d^{-1}$) | **Season*** ($kg\ C\ km^{-2}\ 120\ d^{-1}$) | |
| NPP | (i) | $5.16 \times 10^2$ | $6.19 \times 10^4$ | $6.19 \times 10^4$ |
| | (ii) | $3.21 \times 10^2$ | $3.85 \times 10^4$ | $3.85 \times 10^4$ |
| | (iii) | $8.67 \times 10^2$ | $1.04 \times 10^5$ | $1.04 \times 10^5$ |
| NECB | (i) | $5.16 \times 10^2$ | $6.19 \times 10^4$ | $6.19 \times 10^4$ |
| | (ii) | $3.21 \times 10^2$ | $3.85 \times 10^4$ | $3.22 \times 10^4$ |
| | (iii) | $8.66 \times 10^2$ | $1.04 \times 10^5$ | $9.44 \times 10^4$ |

*Estimated assuming a growing season of 120 days.

### S.5. On the timeframe in which animal-driven carbon effects could materialize

Our analysis illustrates that trophic rewilding pumas and guanacos to Patagonian grassland ecosystems has potential to increase carbon capture and storage and thus to meaningfully add to the portfolio of nature-based solutions. However, one of the uncertainties in using large animal trophic rewilding to increase ecosystem carbon capture and storage in grassland ecosystems is the time frame over which there will be measurable effects on ecosystem carbon storage. There are three published studies that can provide insights to answer this. In Konza Prairie, USA, researchers measured increased grassland primary productivity and soil carbon flux 8–10 years after American bison reintroduction (Johnson & Matchett, 2001). More recently, evidence of increased soil carbon was detected 2–6 years after reintroduction of Exmoor Ponies, European Bison, and Aurochs to a shrub-steppe grassland in Czech Republic (Kaštovská et al., 2024). Conversely, excluding native grazers—e.g., Ibex, Yak, and Bharal—from plots in a Himalayan alpine grassland led to differences in soil C dynamics between fenced and unfenced plots within 4–6 years of the experiment's start—and these differences persisted 8 years later (Naidu et al. 2022). Using animal exclosures in the Patagonia NP rewilding area to compare inside vs. outside conditions could be a fruitful avenue for future research.

### S.6. Local sensitivity analyses of model estimates to parameter values

Here, we present additional tables presenting insights into the sensitivity of model estimates of ecosystem gross (GPP) and net (NPP) primary productivity, as well as carbon sink strength (aka net ecosystem carbon balance, NECB), to the values of key parameters for each modeling scenario. We focus on four key parameters: the soil carbon loss rate ($q_S$; Table S.2), the plant recycling rate ($r_P$; Table S.3), the plant respiration rate (contributing to $\delta$; Table S.4), the herbivore uptake rate ($a_H$; Table S.5), the herbivore recycling rate ($r_H$; Table S.6) and the predator uptake rate ($a_R$; Table S.7). Of these, $q_S$, $r_P$, and the plant respiration rate appear in all three modeling scenarios, whereas $a_H$ and $r_H$ appear in scenarios (2) and (3), and $a_R$ appears only in scenario (3). The sensitivity analyses reveal that the model estimates can vary from −45% below up to +122% above the median value but is often much lower. However, it also shows that in some cases (Table S.2, S.4, and S.5), parameter variation can lead to predictions that the ecosystem can flip from being a carbon sink to a source (i.e., negative lower bounds for yearly NECB estimates).

Table S.5. Median and interquartile range for daily, seasonal, and yearly model estimates of gross primary productivity (GPP), net primary productivity (NPP), and net ecosystem carbon balance (i.e., carbon sink strength; NECB), when the value of soil carbon loss rate (parameter $q_S$) varies 30% above and below the value reported in Table 2. Note that (i) estimates are reported on a per-km$^2$ scale and (ii) NECB estimates account for guanaco methane ($CH_4$) emissions and, at the yearly time scale, for guanaco and puma respiration outside the growing season.

| | | **Daily** (kg C km$^{-2}$ d$^{-1}$) | | **Season** (kg C km$^{-2}$)* | | **Yearly** (kg C km$^{-2}$ y$^{-1}$)* | |
|---|---|---|---|---|---|---|---|
| | **Scenario** | **Median** | **Range†** | **Median** | **Range†** | **Median** | **Range†** |
| GPP | No animal | $6.44 \times 10^2$ | $4.51 \times 10^2$–$8.35 \times 10^2$ | $7.73 \times 10^4$ | $5.41 \times 10^4$–$1.00 \times 10^5$ | $7.73 \times 10^4$ | $5.41 \times 10^4$–$1.00 \times 10^5$ |
| | Guanaco | $4.12 \times 10^2$ | $3.17 \times 10^2$–$5.89 \times 10^2$ | $4.95 \times 10^4$ | $3.81 \times 10^4$–$7.07 \times 10^4$ | $4.95 \times 10^4$ | $3.81 \times 10^4$–$7.07 \times 10^4$ |
| | Guanaco-Puma | $1.11 \times 10^3$ | $7.81 \times 10^2$–$1.45 \times 10^3$ | $1.33 \times 10^5$ | $9.38 \times 10^4$–$1.74 \times 10^5$ | $1.33 \times 10^5$ | $9.38 \times 10^4$–$1.74 \times 10^5$ |
| NPP | No animal | $5.17 \times 10^2$ | $3.62 \times 10^2$–$6.70 \times 10^2$ | $6.20 \times 10^4$ | $4.34 \times 10^4$–$8.04 \times 10^4$ | $6.20 \times 10^4$ | $4.34 \times 10^4$–$8.04 \times 10^4$ |
| | Guanaco | $3.21 \times 10^2$ | $2.26 \times 10^2$–$4.98 \times 10^2$ | $3.85 \times 10^4$ | $2.71 \times 10^4$–$5.98 \times 10^4$ | $3.85 \times 10^4$ | $2.71 \times 10^4$–$5.98 \times 10^4$ |
| | Guanaco-Puma | $8.66 \times 10^2$ | $6.09 \times 10^2$–$1.13 \times 10^3$ | $1.04 \times 10^5$ | $7.30 \times 10^4$–$1.35 \times 10^5$ | $1.04 \times 10^5$ | $7.30 \times 10^4$–$1.35 \times 10^5$ |
| NECB | No animal | $5.17 \times 10^2$ | $3.62 \times 10^2$–$6.60 \times 10^2$ | $6.20 \times 10^4$ | $4.34 \times 10^4$–$8.04 \times 10^4$ | $6.20 \times 10^4$ | $4.34 \times 10^4$–$8.04 \times 10^4$ |
| | Guanaco | $3.20 \times 10^2$ | $2.39 \times 10^2$–$4.71 \times 10^2$ | $3.84 \times 10^4$ | $2.87 \times 10^4$–$5.66 \times 10^4$ | $3.24 \times 10^4$ | $-1.94 \times 10^5$–$1.54 \times 10^5$ |
| | Guanaco-Puma | $8.65 \times 10^2$ | $6.07 \times 10^2$–$1.13 \times 10^3$ | $1.04 \times 10^5$ | $7.29 \times 10^4$–$1.35 \times 10^5$ | $9.44 \times 10^4$ | $8.98 \times 10^4$–$9.91 \times 10^4$ |

* Assuming a growing season of 120 days.
† Minimum-Maximum.

Table S.6. Median and interquartile range for daily and seasonal model estimates of gross primary productivity (GPP), net primary productivity (NPP), and net ecosystem carbon balance (i.e., carbon sink strength; NECB), when the value of plant recycling rate (parameter $r_P$) varies 30% above and below the value reported in Table 2. Note that (i) estimates are reported on a per-km$^2$ scale and (ii) NECB estimates account for guanaco methane ($CH_4$) emissions and, at the yearly time scale, for guanaco and puma respiration outside the growing season.

| | | **Daily** (kg C km$^{-2}$ d$^{-1}$) | | **Season** (kg C km$^{-2}$)$^{*}$ | | **Yearly** (kg C km$^{-2}$ y$^{-1}$)$^{*}$ | |
|---|---|---|---|---|---|---|---|
| | **Scenario** | **Median** | **Range**$^{\dagger}$ | **Median** | **Range**$^{\dagger}$ | **Median** | **Range**$^{\dagger}$ |
| GPP | No animal | $6.44 \times 10^2$ | $4.88 \times 10^2$–$7.97 \times 10^2$ | $7.73 \times 10^4$ | $5.86 \times 10^4$–$9.57 \times 10^4$ | $7.73 \times 10^4$ | $5.86 \times 10^4$–$9.57 \times 10^4$ |
| | Guanaco | $4.09 \times 10^2$ | $2.64 \times 10^2$–$5.66 \times 10^2$ | $4.91 \times 10^4$ | $3.17 \times 10^4$–$6.79 \times 10^4$ | $4.91 \times 10^4$ | $3.17 \times 10^4$–$6.79 \times 10^4$ |
| | Guanaco-Puma | $1.11 \times 10^3$ | $8.64 \times 10^2$–$1.36 \times 10^3$ | $1.34 \times 10^5$ | $1.04 \times 10^5$–$1.64 \times 10^5$ | $1.34 \times 10^5$ | $1.04 \times 10^5$–$1.64 \times 10^5$ |
| NPP | No animal | $5.17 \times 10^2$ | $3.61 \times 10^2$–$6.70 \times 10^2$ | $6.21 \times 10^4$ | $4.34 \times 10^4$–$8.05 \times 10^4$ | $6.21 \times 10^4$ | $4.34 \times 10^4$–$8.05 \times 10^4$ |
| | Guanaco | $3.18 \times 10^2$ | $1.80 \times 10^2$–$4.69 \times 10^2$ | $3.82 \times 10^4$ | $2.16 \times 10^4$–$5.63 \times 10^4$ | $3.82 \times 10^4$ | $2.16 \times 10^4$–$5.63 \times 10^4$ |
| | Guanaco-Puma | $8.69 \times 10^2$ | $6.17 \times 10^2$–$1.12 \times 10^3$ | $1.04 \times 10^5$ | $7.40 \times 10^4$–$1.34 \times 10^5$ | $1.04 \times 10^5$ | $7.40 \times 10^4$–$1.34 \times 10^5$ |
| NECB | No animal | $5.17 \times 10^2$ | $3.61 \times 10^2$–$6.70 \times 10^2$ | $6.21 \times 10^4$ | $4.34 \times 10^4$–$8.05 \times 10^4$ | $6.21 \times 10^4$ | $4.34 \times 10^4$–$8.05 \times 10^4$ |
| | Guanaco | $3.18 \times 10^2$ | $1.84 \times 10^2$–$4.64 \times 10^2$ | $3.81 \times 10^4$ | $2.20 \times 10^4$–$5.57 \times 10^4$ | $3.27 \times 10^4$ | $6.92 \times 10^3$–$5.27 \times 10^4$ |
| | Guanaco-Puma | $8.68 \times 10^2$ | $6.16 \times 10^2$–$1.12 \times 10^3$ | $1.04 \times 10^5$ | $7.39 \times 10^4$–$1.34 \times 10^5$ | $9.46 \times 10^4$ | $5.53 \times 10^4$–$1.29 \times 10^5$ |

$^{*}$ Assuming a growing season of 120 days.
$^{\dagger}$ Minimum-Maximum.

Table S.7. Median and interquartile range for daily and seasonal model estimates of gross primary productivity (GPP), net primary productivity (NPP), and net ecosystem carbon balance (i.e., carbon sink strength; NECB), when the value of plant respiration rate (contributing to parameter $\delta$) varies 30% above and below the value reported in Table 2. Note that (i) estimates are reported on a per-$km^2$ scale and (ii) NECB estimates account for guanaco methane ($CH_4$) emissions and, at the yearly time scale, for guanaco and puma respiration outside the growing season.

| | | **Daily** (kg C $km^{-2}$ $d^{-1}$) | | **Season** (kg C $km^{-2}$)* | | **Yearly** (kg C $km^{-2}$ $y^{-1}$)* | |
|---|---|---|---|---|---|---|---|
| | **Scenario** | **Median** | **Range**† | **Median** | **Range**† | **Median** | **Range**† |
| GPP | No animal | $6.41 \times 10^2$ | $6.05 \times 10^2$–$6.81 \times 10^2$ | $7.70 \times 10^4$ | $7.26 \times 10^4$–$8.17 \times 10^4$ | $7.70 \times 10^4$ | $7.26 \times 10^4$–$8.17 \times 10^4$ |
| | Guanaco | $4.14 \times 10^2$ | $3.87 \times 10^2$–$4.42 \times 10^2$ | $4.96 \times 10^4$ | $4.64 \times 10^4$–$5.31 \times 10^4$ | $4.96 \times 10^4$ | $4.64 \times 10^4$–$5.31 \times 10^4$ |
| | Guanaco-Puma | $1.11 \times 10^3$ | $1.04 \times 10^3$–$1.19 \times 10^3$ | $1.33 \times 10^5$ | $1.25 \times 10^5$–$1.43 \times 10^5$ | $1.33 \times 10^5$ | $1.25 \times 10^5$–$1.43 \times 10^5$ |
| NPP | No animal | $5.16 \times 10^2$ | $5.16 \times 10^2$–$5.16 \times 10^2$ | $6.19 \times 10^4$ | $6.19 \times 10^4$–$6.19 \times 10^4$ | $6.19 \times 10^4$ | $6.19 \times 10^4$–$6.19 \times 10^4$ |
| | Guanaco | $3.23 \times 10^2$ | $2.75 \times 10^2$–$3.74 \times 10^2$ | $3.88 \times 10^4$ | $3.30 \times 10^4$–$4.48 \times 10^4$ | $3.88 \times 10^4$ | $3.30 \times 10^4$–$4.48 \times 10^4$ |
| | Guanaco-Puma | $8.67 \times 10^2$ | $8.66 \times 10^2$–$8.68 \times 10^2$ | $1.04 \times 10^5$ | $1.04 \times 10^5$–$1.04 \times 10^5$ | $1.04 \times 10^5$ | $1.04 \times 10^5$–$1.04 \times 10^5$ |
| NECB | No animal | $5.16 \times 10^2$ | $5.16 \times 10^2$–$5.16 \times 10^2$ | $6.19 \times 10^4$ | $6.19 \times 10^4$–$6.19 \times 10^4$ | $6.19 \times 10^4$ | $6.19 \times 10^4$–$6.19 \times 10^4$ |
| | Guanaco | $3.33 \times 10^2$ | $2.78 \times 10^2$–$3.69 \times 10^2$ | $3.87 \times 10^4$ | $3.34 \times 10^4$–$4.42 \times 10^4$ | $3.09 \times 10^4$ | $-3.27 \times 10^3$–$6.32 \times 10^4$ |
| | Guanaco-Puma | $8.66 \times 10^2$ | $8.65 \times 10^2$–$8.68 \times 10^2$ | $1.04 \times 10^5$ | $1.04 \times 10^5$–$1.04 \times 10^5$ | $9.46 \times 10^4$ | $9.86 \times 10^4$–$1.00 \times 10^5$ |

* Assuming a growing season of 120 days.
† Minimum-Maximum.

Table S.8. Median and interquartile range for daily and seasonal model estimates of gross primary productivity (GPP), net primary productivity (NPP), and net ecosystem carbon balance (i.e., carbon sink strength; NECB), when the value of herbivore uptake rate (parameter $a_H$) varies 30% above and below the value reported in Table 2. Note that (i) only two modeling scenarios are reported here as parameter $a_H$ is set to 0 in scenario (1), (ii) estimates are reported on a per-km$^2$ scale, and (ii) NECB estimates account for guanaco methane ($CH_4$) emissions and, at the yearly time scale, for guanaco and puma respiration outside the growing season.

| | | **Daily** (kg C km$^{-2}$ d$^{-1}$) | | **Season** (kg C km$^{-2}$ 120 d$^{-1}$)* | | **Yearly** (kg C km$^{-2}$ y$^{-1}$)* | |
|---|---|---|---|---|---|---|---|
| | **Scenario** | **Median** | **Range**† | **Median** | **Range**† | **Median** | **Range**† |
| GPP | Guanaco | $4.11 \times 10^2$ | $2.45 \times 10^2$–$8.42 \times 10^2$ | $4.93 \times 10^4$ | $2.93 \times 10^4$–$1.01 \times 10^5$ | $4.93 \times 10^4$ | $2.93 \times 10^4$–$1.01 \times 10^5$ |
| | Guanaco-Puma | $1.11 \times 10^3$ | $1.10 \times 10^3$–$1.12 \times 10^3$ | $1.34 \times 10^5$ | $1.33 \times 10^5$–$1.35 \times 10^5$ | $1.34 \times 10^5$ | $1.34 \times 10^5$–$1.35 \times 10^5$ |
| NPP | Guanaco | $3.20 \times 10^2$ | $1.74 \times 10^2$–$7.11 \times 10^2$ | $3.84 \times 10^4$ | $2.09 \times 10^4$–$8.54 \times 10^4$ | $3.84 \times 10^4$ | $2.09 \times 10^4$–$8.54 \times 10^4$ |
| | Guanaco-Puma | $8.67 \times 10^2$ | $8.59 \times 10^2$–$8.75 \times 10^2$ | $1.04 \times 10^5$ | $1.03 \times 10^5$–$1.05 \times 10^5$ | $1.04 \times 10^5$ | $1.03 \times 10^5$–$1.05 \times 10^5$ |
| NECB | Guanaco | $3.19 \times 10^2$ | $1.84 \times 10^2$–$6.73 \times 10^2$ | $3.83 \times 10^4$ | $2.21 \times 10^4$–$8.08 \times 10^4$ | $3.32 \times 10^4$ | $-2.78 \times 10^5$–$1.19 \times 10^5$ |
| | Guanaco-Puma | $8.66 \times 10^2$ | $8.58 \times 10^2$–$8.75 \times 10^2$ | $1.04 \times 10^5$ | $1.03 \times 10^5$–$1.05 \times 10^5$ | $9.54 \times 10^4$ | $6.83 \times 10^4$–$1.20 \times 10^5$ |

* Assuming a growing season of 120 days.
† Minimum-Maximum.

Table S.9. Median and interquartile range for daily and seasonal model estimates of gross primary productivity (GPP), net primary productivity (NPP), and net ecosystem carbon balance (i.e., carbon sink strength; NECB), when the value of herbivore recycling rate (parameter $r_H$) varies 30% above and below the value reported in Table 2. Note that (i) only two modeling scenarios are reported here as parameter $r_H$ is set to 0 in scenario (1), (ii) estimates are reported on a per-$km^2$ scale, and (ii) NECB estimates account for guanaco methane ($CH_4$) emissions and, at the yearly time scale, for guanaco and puma respiration outside the growing season.

| | | **Daily** (kg C $km^{-2}$ $d^{-1}$) | | **Season** (kg C $km^{-2}$ 120 $d^{-1}$)* | | **Yearly** (kg C $km^{-2}$ $y^{-1}$)* | |
|---|---|---|---|---|---|---|---|
| | **Scenario** | **Median** | **Range**† | **Median** | **Range**† | **Median** | **Range**† |
| GPP | Guanaco | $4.19 \times 10^2$ | $2.03 \times 10^2$–$6.97 \times 10^2$ | $5.03 \times 10^4$ | $2.43 \times 10^4$–$8.36 \times 10^4$ | $5.03 \times 10^4$ | $2.43 \times 10^4$–$8.36 \times 10^4$ |
| | Guanaco-Puma | $1.11 \times 10^3$ | $1.11 \times 10^3$–$1.11 \times 10^3$ | $1.34 \times 10^5$ | $1.34 \times 10^5$–$1.34 \times 10^5$ | $1.34 \times 10^5$ | $1.34 \times 10^5$–$1.34 \times 10^5$ |
| NPP | Guanaco | $3.27 \times 10^2$ | $1.39 \times 10^2$–$5.78 \times 10^2$ | $3.92 \times 10^4$ | $1.66 \times 10^4$–$6.93 \times 10^4$ | $3.92 \times 10^4$ | $1.66 \times 10^4$–$6.93 \times 10^4$ |
| | Guanaco-Puma | $8.67 \times 10^2$ | $8.67 \times 10^2$–$8.67 \times 10^2$ | $1.04 \times 10^5$ | $1.04 \times 10^5$–$1.04 \times 10^5$ | $1.04 \times 10^5$ | $1.04 \times 10^5$–$1.04 \times 10^5$ |
| NECB | Guanaco | $3.26 \times 10^2$ | $1.56 \times 10^2$–$5.59 \times 10^2$ | $3.91 \times 10^4$ | $1.88 \times 10^4$–$6.71 \times 10^4$ | $2.84 \times 10^4$ | $-1.10 \times 10^5$–$1.83 \times 10^5$ |
| | Guanaco-Puma | $8.66 \times 10^2$ | $8.66 \times 10^2$–$8.67 \times 10^2$ | $1.04 \times 10^5$ | $1.04 \times 10^5$–$1.04 \times 10^5$ | $9.47 \times 10^4$ | $6.79 \times 10^4$–$1.21 \times 10^5$ |

* Assuming a growing season of 120 days.
† Minimum-Maximum.

Table S.10. Median and interquartile range for daily and seasonal model estimates of gross primary productivity (GPP), net primary productivity (NPP), and net ecosystem carbon balance (i.e., carbon sink strength; NECB), when the value of predator uptake rate (parameter $a_R$) varies 30% above and below the value reported in Table 2. Note that only one modeling scenario is reported here as parameter $a_R$ is set to 0 in scenarios (1) and (2). Note also that (i) estimates are reported on a per-km$^2$ scale and (ii) NECB estimates account for guanaco methane ($CH_4$) emissions and, at the yearly time scale, for guanaco and puma respiration outside the growing season.

| | **Daily** (kg C km$^{-2}$ d$^{-1}$) | | **Season** (kg C km$^{-2}$)$^{*}$ | | **Yearly** (kg C km$^{-2}$ y$^{-1}$)$^{*}$ | |
|---|---|---|---|---|---|---|
| | **Median** | **Range**$^{†}$ | **Median** | **Range**$^{†}$ | **Median** | **Range**$^{†}$ |
| GPP | 1.11 × 10$^3$ | 1.11 × 10$^3$–1.13 × 10$^3$ | 1.34 × 10$^5$ | 1.33 × 10$^5$–1.35 × 10$^5$ | 1.34 × 10$^5$ | 1.33 × 10$^5$–1.35 × 10$^5$ |
| NPP | 8.67 × 10$^2$ | 8.61 × 10$^2$–8.79 × 10$^2$ | 1.04 × 10$^5$ | 1.03 × 10$^5$–1.05 × 10$^5$ | 1.04 × 10$^5$ | 1.03 × 10$^5$–1.05 × 10$^5$ |
| NECB | 8.66 × 10$^2$ | 8.61 × 10$^2$–8.77 × 10$^2$ | 1.04 × 10$^5$ | 1.03 × 10$^5$–1.05 × 10$^5$ | 9.44 × 10$^4$ | 9.09 × 10$^4$–9.61 × 10$^4$ |

$^{*}$ Assuming a growing season of 120 days.
$^{†}$ Minimum-Maximum.

### S.7. Supplementary Tables

Table S.11. Parameter values used in the model, showing the original ("Raw value") parameter values sourced from the literature, the parameter values used in the final, final model ("Adjusted value"), and the factor by which the parameter was varied in the final model ("Adjustment factor"). For details and rationale on why parameters were adjusted, see main text and Supporting Code document in the online code repository.

| Scenario | Description | Parameter | Raw value | Adjusted value | Adjustment factor | Units | Notes | Source |
|---|---|---|---|---|---|---|---|---|
| i, ii, iii | N inorganic inputs | $I$ | $9.30 \times 10^{-7}$ | $9.30 \times 10^{-7}$ | | kg N $(m^2\ d)^{-1}$ | | Godoy et al., 2003 |
| i, ii, iii | Plant uptake rate | $a_P$ | $2.50 \times 10^{-1}$ | $2.50 \times 10^{-1}$ | | % $d^{-1}$ | | Gherardi et al., 2013 |
| ii | Herbivore uptake rate | $a_H$ | $1.33 \times 10^{-3}$ | $1.33 \times 10^{-1}$ | $1.00 \times 10^{2}$ | % N (kg Guanaco N $d)^{-1}$ | | Meyer et al., 2010 |
| iii | Herbivore uptake rate | $a_H$ | $1.07 \times 10^{-3}$ | $1.07 \times 10^{-1}$ | $1.00 \times 10^{2}$ | % N (kg Guanaco N $d)^{-1}$ | Adjusted to 75% of herbivore-only value to account for reduction due to predation | Meyer et al., 2010 |
| iii | Predator uptake rate | $a_R$ | $8.97 \times 10^{-7}$ | $8.97 \times 10^{-2}$ | $1.00 \times 10^{5}$ | % N (kg Puma N $d)^{-1}$ | | Bank et al., 2002, Cristescu et al. (2022) |
| i | Plant respiration rate | *contributes to* $\delta$ | $1.22 \times 10^{-3}$ | $1.22 \times 10^{-3}$ | | $d^{-1}$ | | Peri et al., 2024 |
| ii, iii | Plant respiration rate | *contributes to* $\delta$ | $1.09 \times 10^{-3}$ | $1.09 \times 10^{-3}$ | | $d^{-1}$ | Adjusted for herbivory as 89% of no herbivore value | Peri et al., 2024 |
| ii, iii | Herbivore respiration rate | *contributes to* $\pi$ | $2.42 \times 10^{-2}$ | $2.42 \times 10^{-2}$ | | $d^{-1}$ | Box 1. in supplemental of Schmitz et al. 2023 | Schmitz et al., 2023 |
| iii | Predator respiration rate | *contributes to* $\tau$ | $8.58 \times 10^{-3}$ | $8.58 \times 10^{-3}$ | | $d^{-1}$ | Box 1. in supplemental of Schmitz et al. 2023 | Laundré, 2005; Schmitz et al., 2023 |
| i | Plant recycling rate | $r_P$ | $4.97 \times 10^{-6}$ | $4.97 \times 10^{-3}$ | $1.00 \times 10^{3}$ | $d^{-1}$ | | Campanella & Bertiller, 2008 |

| | | | | | | | | |
|---|---|---|---|---|---|---|---|---|
| ii, iii | Plant recycling rate | $r_P$ | $3.77 \times 10^{-6}$ | $3.77 \times 10^{-3}$ | $1.00 \times 10^{3}$ | $d^{-1}$ | | Carrera et al., 2008 |
| ii | Herbivore recycling rate | $r_H$ | $1.64 \times 10^{-3}$ | $5.34 \times 10^{-4}$ | $3.25 \times 10^{-1}$ | $d^{-1}$ | Estimated as 81% of value used for herbivore uptake rate based on source | Meyer et al., 2010 |
| iii | Herbivore recycling rate | $r_H$ | $1.15 \times 10^{-3}$ | $1.15 \times 10^{-3}$ | | $d^{-1}$ | Estimated as 70% of the value without predator based on source | Meyer et al., 2010 |
| iii | Predator recycling rate | $r_R$ | $4.30 \times 10^{-8}$ | $4.30 \times 10^{-5}$ | $1.00 \times 10^{3}$ | $d^{-1}$ | | Bank et al., 2002 |
| i, ii, iii | Plant C:N | $\alpha$ | 27.0 | 27.0 | | unitless | Estimated from data in Table 1 | Perez-Quezada et al., 2022 |
| i, ii, iii | Animal C:N | $\beta$ | 3.29 | 3.29 | | unitless | Estimated as 4.75% of predator uptake rate based on source | Franke & Weniger, 1958 |
| i | Soil C leaching rate | $q_S$ | $2.59 \times 10^{-4}$ | $2.59 \times 10^{-2}$ | $1.00 \times 10^{2}$ | kg N $(m^2\ d)^{-1}$ | | Carbonell-Silletta et al., 2022 |
| ii | Soil C leaching rate | $q_S$ | $2.59 \times 10^{-4}$ | $2.08 \times 10^{-2}$ | $1.00 \times 10^{2}$ | kg N $(m^2\ d)^{-1}$ | Adjusted to 80% of no-animal value before calibration to account for trampling | Carbonell-Silletta et al., 2022 |
| iii | Soil C leaching rate | $q_S$ | $2.59 \times 10^{-4}$ | $5.57 \times 10^{-2}$ | $2.69 \times 10^{2}$ | kg N $(m^2\ d)^{-1}$ | Adjusted to 80% of no-animal value before calibration to account for trampling | Carbonell-Silletta et al., 2022 |
| i, ii, iii | Soil N leaching rate | $k$ | $3.5 \times 10^{-8}$ | $3.50 \times 10^{-4}$ | $1.00 \times 10^{4}$ | $d^{-1}$ | | Yahdjian & Sala, 2010 |

S.8.

### S.9. References

Bank, M. S., Sarno, R. J., Campbell, N. K., & Franklin, W. L. (2002). Predation of guanacos (Lama guanicoe) by southernmost mountain lions (Puma concolor) during a historically severe winter in Torres del Paine National Park, Chile. *Journal of Zoology*, *258*(2), 215–222. https://doi.org/10.1017/S0952836902001334

Campanella, M. V., & Bertiller, M. B. (2008). Plant phenology, leaf traits and leaf litterfall of contrasting life forms in the arid Patagonian Monte, Argentina. *Journal of Vegetation Science*, *19*(1), 75–85. https://doi.org/10.3170/2007-8-18333

Carbonell-Silletta, L., Cavallaro, A., Pereyra, D. A., Askenazi, J. O., Goldstein, G., Scholz, F. G., & Bucci, S. J. (2022). Soil respiration and N-mineralization processes in the Patagonian steppe are more responsive to fertilization than to experimental precipitation increase. *Plant and Soil*, *479*(1), 405–422. https://doi.org/10.1007/s11104-022-05531-0

Carrera, A. L., Bertiller, M. B., & Larreguy, C. (2008). Leaf litterfall, fine-root production, and decomposition in shrublands with different canopy structure induced by grazing in the Patagonian Monte, Argentina. *Plant and Soil*, *311*(1–2), 39–50. https://doi.org/10.1007/s11104-008-9655-8

Chapin, F. S., Woodwell, G. M., Randerson, J. T., Rastetter, E. B., Lovett, G. M., Baldocchi, D. D., Clark, D. A., Harmon, M. E., Schimel, D. S., Valentini, R., Wirth, C., Aber, J. D., Cole, J. J., Goulden, M. L., Harden, J. W., Heimann, M., Howarth, R. W., Matson, P. A., McGuire, A. D., … Schulze, E. D. (2006). Reconciling carbon-cycle concepts, terminology, and methods. *Ecosystems*, *9*(7), 1041–1050. https://doi.org/10.1007/s10021-005-0105-7

Chapin III, F. S., Matson, P. A., & Vitousek, P. M. (2002). *Principles of Terrestrial Ecosystem Ecology* (2nd ed.). Springer New York, NY. https://doi.org/10.1007/978-1-4419-9504-9

Cristescu, B., Elbroch, M.L., Dellinger, J.A., Binder, W., Wilmers, C.C. & Wittmer, H.U. (2022). Kill rates and associated ecological factors for an apex predator. *Mammalian Biology,* 102, 291–305.

Franke, E.-R., & Weniger, J. H. (1958). Der Stickstoff-, Kohlenstoff- und Energiegehalt des Fleisches und der Wärmewert des Fettes verschiedener Nutztierarten. *Archiv Für Tierernaehrung*, *8*(1–6), 81–94. https://doi.org/10.1080/17450395809424770

Gherardi, L. A., Sala, O. E., & Yahdjian, L. (2013). Preference for different inorganic nitrogen forms among plant functional types and species of the Patagonian steppe. *Oecologia*, *173*(3), 1075–1081. https://doi.org/10.1007/s00442-013-2687-7

Godoy, R., Paulino, L., Oyarzún, C., & Boeckx, P. (2003). ATMOSPHERIC N DEPOSITION IN CENTRAL AND SOUTHERN CHILE: AN OVERVIEW. *Gayana. Botánica*, *60*(1), 47–53. https://doi.org/10.4067/S0717-66432003000100008

Johnson, L.C., & Matchett, J.R. (2001). Fire and grazing regulate belowground process in tallgrass prairie. *Ecolog,y* 82, 3377-3389.

Kaštovská, E., Mastný, J., & Konvička, M. (2024). Rewilding by large ungulates contributes to organic carbon storage in soils. *Journal of Environmental Management* 355, 120430. https://doi.org/10.1016/j.jenvman.2024.120430

Laundré, J. W. (2005). Puma Energetics: A Recalculation. *The Journal of Wildlife Management*, *69*(2), 723–732. https://doi.org/10.2193/0022-541X(2005)069%255B0723:PEAR%255D2.0.CO;2

Meyer, K., Hummel, J., & Clauss, M. (2010). The relationship between forage cell wall content and voluntary food intake in mammalian herbivores. *Mammal Review*, *40*(3), 221–245. https://doi.org/10.1111/j.1365-2907.2010.00161.x

Naidu, D.G.T., Roy, S., & Bagchi, S. (2022) Loss of grazing by large mammalian herbivores can destabilize the soil carbon pool. *Proceedings of the National Academy of Science U.S.A.,* 119, 2211317119. https://doi.org/10.1073/pnas.2211317119

Perez-Quezada, J. F., Cano, S., Ibaceta, P., Aguilera-Riquelme, D., Salazar, O., Fuentes, J. P., & Osborne, B. (2022). How do land cover changes affect carbon-nitrogen-phosphorus stocks and the greenhouse gas budget of ecosystems in southern Chile? *Agriculture, Ecosystems & Environment*, *340*, 108153. https://doi.org/10.1016/j.agee.2022.108153

Peri, P. L., Gyenge, J., & Fernández, M. E. (2024). Photosynthetic response to water stress of the main grass species of southern patagonian steppe, Argentina. *Journal of Arid Environments*, *224*, 105219. https://doi.org/10.1016/j.jaridenv.2024.105219

Rizzuto, M., Leroux, S. J., & Schmitz, O. J. (2024). Rewiring the Carbon Cycle: A Theoretical Framework for Animal-Driven Ecosystem Carbon Sequestration. *Journal of Geophysical Research: Biogeosciences*, *129*(4), e2024JG008026. https://doi.org/10.1029/2024JG008026

Schmitz, O. J., Sylvén, M., Atwood, T. B., Bakker, E. S., Berzaghi, F., Brodie, J. F., Cromsigt, J. P. G. M., Davies, A. B., Leroux, S. J., Schepers, F. J., Smith, F. A., Stark, S., Svenning, J.-C., Tilker, A., & Ylänne, H. (2023). Trophic rewilding can expand natural climate solutions. *Nature Climate Change*, 1–10. https://doi.org/10.1038/s41558-023-01631-6

Yahdjian, L., & Sala, O. E. (2010). Size of Precipitation Pulses Controls Nitrogen Transformation and Losses in an Arid Patagonian Ecosystem. *Ecosystems*, *13*(4), 575–585. https://doi.org/10.1007/s10021-010-9341-6
